# Asymptomatic colonization by *Botrytis cinerea* reveals polarized plant-fungus interfaces before widespread disease

**DOI:** 10.64898/2026.10.12.758352

**Authors:** Ines Tguafaiti, Perrine Ferdinand, Adrien Pons, Bastien Malbert, Amélie De Vallée, Christophe Bruel, Mathias Choquer, Nathalie Poussereau, Caroline Bournaud, François-Xavier Gillet

**Author notes:** co-authors (Caroline Bournaud, François-Xavier Gillet).

## Abstract

- *Botrytis cinerea* is a necrotrophic pathogen whose destructive growth is typically studied under conditions that rapidly generate visible disease. Yet symptomless associations are well documented, the fungal and host processes operating before widespread tissue collapse remain poorly resolved.
- We developped an intact *Arabidopsis thaliana* seedling pathosystem that extends the interval between infection establishment and widespread visible disease. Fungal colonization was quantified using qPCR and a constitutively bioluminescent reporter strain, together with longitudinal plant phenotyping, hormone profiling and multiscale fluorescence microscopy.
- Substantial fungal colonization occurred while seedlings remained largely green and continued to grow, whereas robust whole-seedling accumulation of salicylic and jasmonic acids occurred later. During this interval, *B. cinerea* displayed diverse tissue-associated infection morphologies. At individual contact sites, fungal walls showed heterogeneous exposure of chitin-associated epitopes, while host cells exhibited focal plasma-membrane polarization toward fungal contact sites, together with cortical actin reorganization.
- These results show that the interval preceding widespread visible disease is an active and spatially heterogeneous interaction state, in which substantial fungal development and highly organized local host responses are already established. Intact seedlings therefore provide a powerful system for resolving plant–pathogen interactions that are largely obscured in rapid- disease assays.

## Introduction

Botrytis cinerea is a broad-host-range fungal pathogen and the causal agent of grey mould disease in numerous plant species. As a paradigmatic necrotroph, its pathogenicity is associated with the coordinated deployment of plant cell wall-degrading enzymes, phytotoxic metabolites, cell death-inducing proteins and other secreted factors that ultimately promote host tissue collapse and exploitation (Veloso & van Kan, 2018; Bi et al., 2023). This destructive outcome is preceded by an ordered sequence of fungal and host events. In Arabidopsis thaliana, time-resolved transcriptomic analyses have revealed extensive reprogramming of both partners before extensive lesion development, including fungal morphogenesis together with changes in host defence, metabolism and hormone-associated pathways (Windram et al., 2012; Wei et al., 2024). Most experimental pathosystems, however, are designed to favour a rapid and reproducible transition from infection establishment to visible disease, commonly using localized inoculation of detached or intact leaves with relatively concentrated conidial suspensions under high humidity and often with exogenous nutrient supplementation. Under such conditions, fungal germination, penetration and invasive growth occur within the first one to two days and are rapidly followed by extensive mycelial expansion and tissue collapse. Although well suited to quantify disease development, such assays provide limited temporal resolution of the interaction states that precede widespread necrotic growth, even though fungal morphogenesis and host reprogramming are already underway during this interval.

The conditions used to accelerate disease influence fungal developmental behaviour. Conidial density modulates penetration rate, lesion progression and the differentiation of infection structures in B. cinerea (van den Heuvel & Waterreus, 1983). Nutrient availability likewise regulates conidial germination and early infection development (Blakeman, 1975; Doehlemann et al., 2006; Liu et al., 2018). Beyond their nutritional effect, plant-derived components present in inoculation media can directly influence fungal development and virulence (Escaray et al., 2024). In contrast, persistent and systemic symptomless colonization has been observed following seed-borne infection or relatively sparse dry conidial inoculation of young plants in Primula, lettuce and Arabidopsis (Barnes & Shaw, 2002, 2003; Sowley et al., 2010; Shaw et al., 2016). B. cinerea could subsequently be recovered from apparently healthy tissues and from organs that developed after inoculation. Targeted RT-qPCR analysis of fungal pathogenicity-related genes further suggested that fungal activity during symptomless colonization differs from that associated with established necrotic disease (Emmanuel et al., 2018). Microscopy directly revealed fungal growth on and within symptomless lettuce and Arabidopsis tissues (Emmanuel et al., 2023). Thus, the absence of visible symptoms does not imply fungal inactivity. Overall, experimental conditions can strongly shape which interaction states are accessible for analysis, with lower inoculum pressure and limited exogenous nutrient availability potentially extending the interval between infection establishment and widespread visible disease.

The host environment encountered by a pathogen varies with the physiological and developmental state of the host. In Arabidopsis, resistance to B. cinerea differs among tissues and changes with developmental age (Dai et al., 2019). Defence responses are also spatially organized within infected tissues, differing between regions close to and distant from lesions (Mulema & Denby, 2012). Host physiological state can also strongly influence disease progression. In unripe tomato fruit, the transition to destructive disease depends on ripening, and B. cinerea can itself accelerate ripening-associated processes (Silva et al., 2023). Seedling establishment is a particularly dynamic developmental window, during which rapid growth and tissue differentiation coincide with major changes in immune competence. In Arabidopsis, components of basal innate immunity progressively mature during the first days after germination (Zou et al., 2018). Responses to the necrotrophic fungus Alternaria brassicicola also differ markedly between germinating seeds and later seedling stages (Ortega-Cuadros et al., 2022). In the Arabidopsis–B. cinerea interaction itself, a regulator of juvenile development modulates JA signalling and susceptibility to the fungus (Sun et al., 2022). Together, these observations suggest that a seedling is not merely a smaller host, but a rapidly changing and spatially heterogeneous physiological environment in which infection outcome may vary with developmental and tissue context. Studying B. cinerea in this developmental context while extending the interval preceding widespread visible disease may therefore reveal context-dependent fungal behaviours and plant–fungus interfaces that remain poorly explored.

Here, we investigated fungal colonization and plant–fungus interface organization during infection of intact *A. thaliana* seedlings by *B. cinerea*. We established an intact-seedling pathosystem combining low inoculum pressure with the absence of exogenous nutrient supplementation, thereby extending the interval between infection establishment and widespread visible disease while enabling fungal colonization to be quantified over time. We quantified the temporal relationship between fungal accumulation, host phenotypic changes and defence-associated hormone accumulation, examined the distribution of infection-related fungal morphologies across seedling tissues, and characterized local plant–fungus interfaces at the cellular scale. To do so, we combined longitudinal plant phenotyping with two independent quantitative readouts of fungal colonization, hormone profiling and multiscale fluorescence microscopy. This approach provides a quantitative and spatially resolved view of the interaction period preceding widespread visible disease in an intact developing host.

## Results

### Active *B. cinerea* colonization precedes visible disease in *Arabidopsis* seedlings

We established an *Arabidopsis* seedling pathosystem using a low spore load delivered in water to seedlings grown on sucrose-free medium, conditions chosen to limit exogenous nutritional and signalling inputs. This approach was motivated in part by evidence that plant-derived components of conventional inoculation media can alter *B. cinerea* growth and developmental programmes (Escaray *et al*., 2024). To enable sensitive monitoring of fungal progression throughout infection, we generated the Hemera reporter strain from B05.10. A constitutive NanoLuc (nLuc) expression cassette was inserted at an intergenic site between *BcSRP68* and *BcNIA-D*, adjacent to a locus previously used for targeted reporter integration in *B. cinerea* (Schumacher, 2012) - (Figure 1a; Figure S1). Hemera showed no obvious *in vitro* growth defect compared with the parental B05.10 strain (Figure S1). Inoculum deposition and spore germination were monitored in parallel (Figure S2).

**Figure 1.**
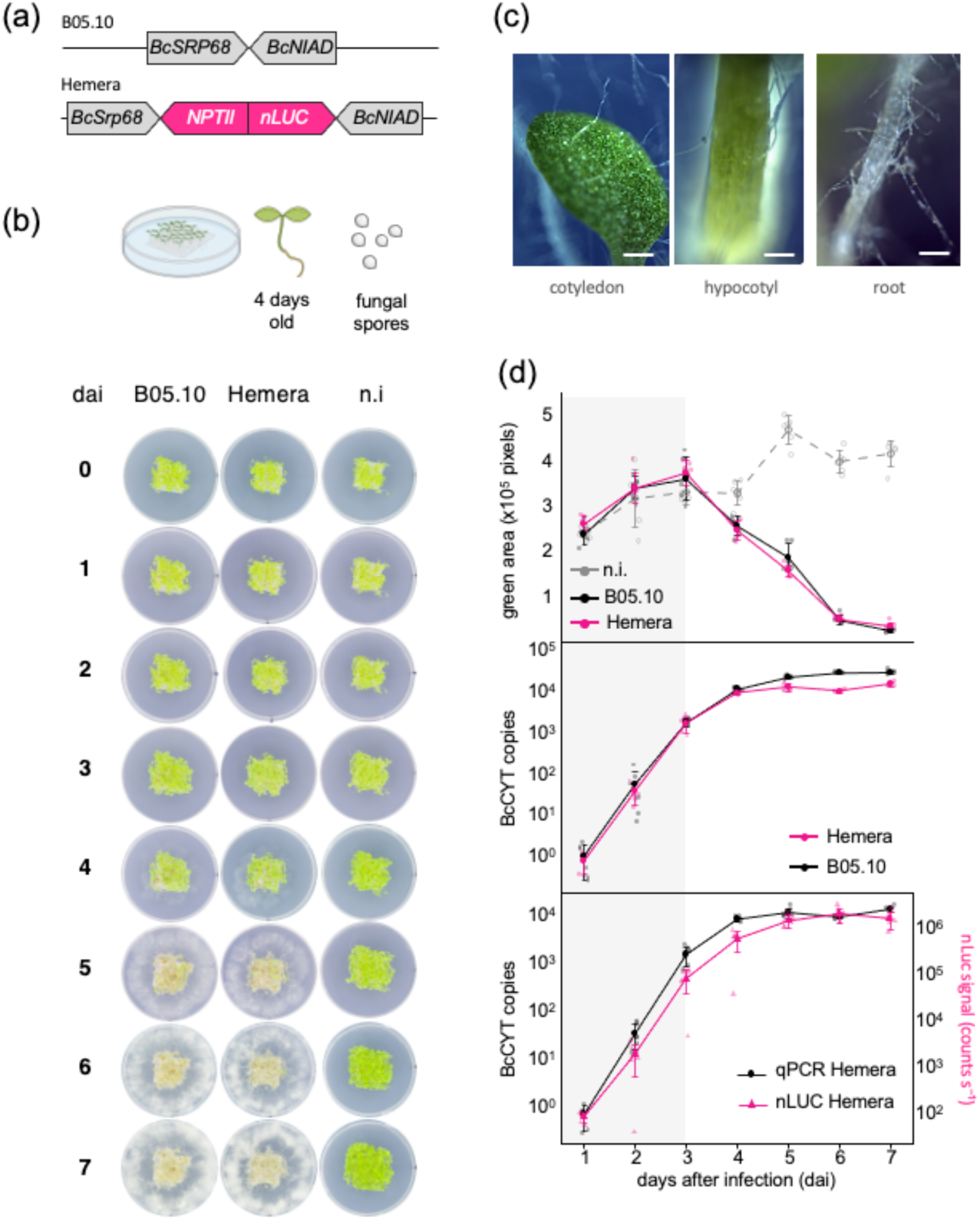
Active *Botrytis cinerea* colonization precedes widespread visible disease in *Arabidopsis* seedlings. **(a)** Generation of the constitutive NanoLuc reporter strain Hemera. A constitutive *nLuc* expression cassette was inserted at an intergenic site between *BcNIAD (Bcin07g01270)* and *BcSRP68 (Bcin07g01260)* to generate the bioluminescent *B. cinerea* Hemera strain. **(b)** Longitudinal monitoring of the Arabidopsis seedling pathotest. Representative images show non-infected seedlings and seedlings inoculated with the parental B05.10 or Hemera strains from 0 to 7 days after inoculation (dai). Fungal growth progressively develops while seedlings remain largely green during the early stages of infection, and later extends onto the surrounding medium before host tissues are fully collapsed. **(c)** Representative stereomicroscope views at 3 dai showing fungal growth in close association with multiple seedling tissues, including the cotyledon, hypocotyl and root. Scale bars, 100 µm. **(d)** Quantification of host development and fungal progression during infection. Seedling development was monitored as green area in non-infected seedlings and seedlings infected with B05.10 or Hemera. Fungal progression was independently quantified by qPCR targeting the mitochondrial *BcCYT* locus and, for Hemera, by NanoLuc activity. The grey shaded region indicates the macroscopically asymptomatic phase, during which fungal progression increases strongly while seedling green area is still maintained or only begins to decline. Points represent biological replicates (n = 6); lines and error bars indicate mean ± SD.

To quantify host-associated fungal progression independently of visible symptoms, we first validated qPCR assays targeting the mitochondrial *BcCYT* and single-copy nuclear *BcPDA* loci on B05.10 strain (Figure S3). Both assays captured highly concordant temporal dynamics of fungal accumulation (r = 0.996, P < 0.001; slope = 1.021), while *BcCYT* yielded approximately 18.8-fold more target copies from the same amount of fungal genomic DNA and consequently improved detection at low fungal loads (Figure S4). *BcCYT* was therefore selected for subsequent infection time courses. We next evaluated NanoLuc bioluminescence as an independent readout of fungal progression. Hemera closely reproduced the temporal infection dynamics of B05.10 (r = 0.997, P < 0.001; slope = 0.936), with no significant strain × day interaction (P = 0.119; Figure S5b). In Hemera-infected seedlings, NanoLuc activity and *BcCYT* accumulation were likewise strongly concordant throughout infection (r = 0.985, P < 0.001; slope = 1.017), with no significant time-dependent change in their relative offset (P = 0.321; Figure S5c). These analyses therefore validated NanoLuc activity as a complementary quantitative proxy for fungal progression in this pathosystem.

Longitudinal imaging of the pathotest showed that infected seedlings remained largely green during the early stages of infection, while fungal growth progressively became visible and subsequently extended onto the surrounding medium (Figure 1b). At 3 dai, stereomicroscopy revealed extensive fungal growth in close association with several seedling tissues, including cotyledons, hypocotyls and roots, although much of the host remained green (Figure 1c ; Figure S6). Quantitative analysis of seedling green area using GreenDayZ (Tguafaiti et al., 2026) confirmed this temporal separation between fungal colonization and visible host decline: seedling green area continued to increase until 3 days after inoculation (dai), while fungal DNA had already increased by approximately three orders of magnitude (Figure 1d). Green area declined thereafter, marking the onset of visible disease. Thus, substantial host-associated fungal progression preceded the widespread loss of green host tissue.

Independent pathotests showed some variation in the exact timing of fungal progression, but the overall infection trajectory remained reproducible (Figure S5a). Most importantly, fungal progression became substantial while seedling green area was still increasing, and infected seedlings remained largely green despite visible fungal growth across several host tissues (Figure 1). We therefore refer to this whole-seedling interval, during which host-associated fungal progression occurs before widespread visible disease, as the macroscopically asymptomatic phase.

### Increasing spore load shortens the pre-expansion phase without altering subsequent fungal expansion

The reproducible infection trajectories suggested that fungal progression could be separated into a prolonged pre-expansion period, a phase of rapid increase and a later plateau. We therefore asked which part of this trajectory was affected by the initial inoculum (Figure 2). An empirically defined NanoLuc activity threshold of 10⁵ was used as an operational marker of the transition to rapid fungal expansion. Increasing the inoculum from 5 to 25 or 500 spores progressively shortened the time required to reach this threshold (Figure 2a,c). By contrast, when infection kinetics were aligned to threshold crossing, the subsequent increase in NanoLuc activity was similar between the 25- and 500-spore inocula (Figure 2b; Figure S7). At the lowest inoculum, NanoLuc activity remained low for several days before increasing sharply, resulting in a markedly prolonged pre-expansion phase (Figure 2a). Thus, spore load primarily affected the duration preceding rapid fungal expansion rather than the kinetics observed once this transition had been reached.

**Figure 2.**
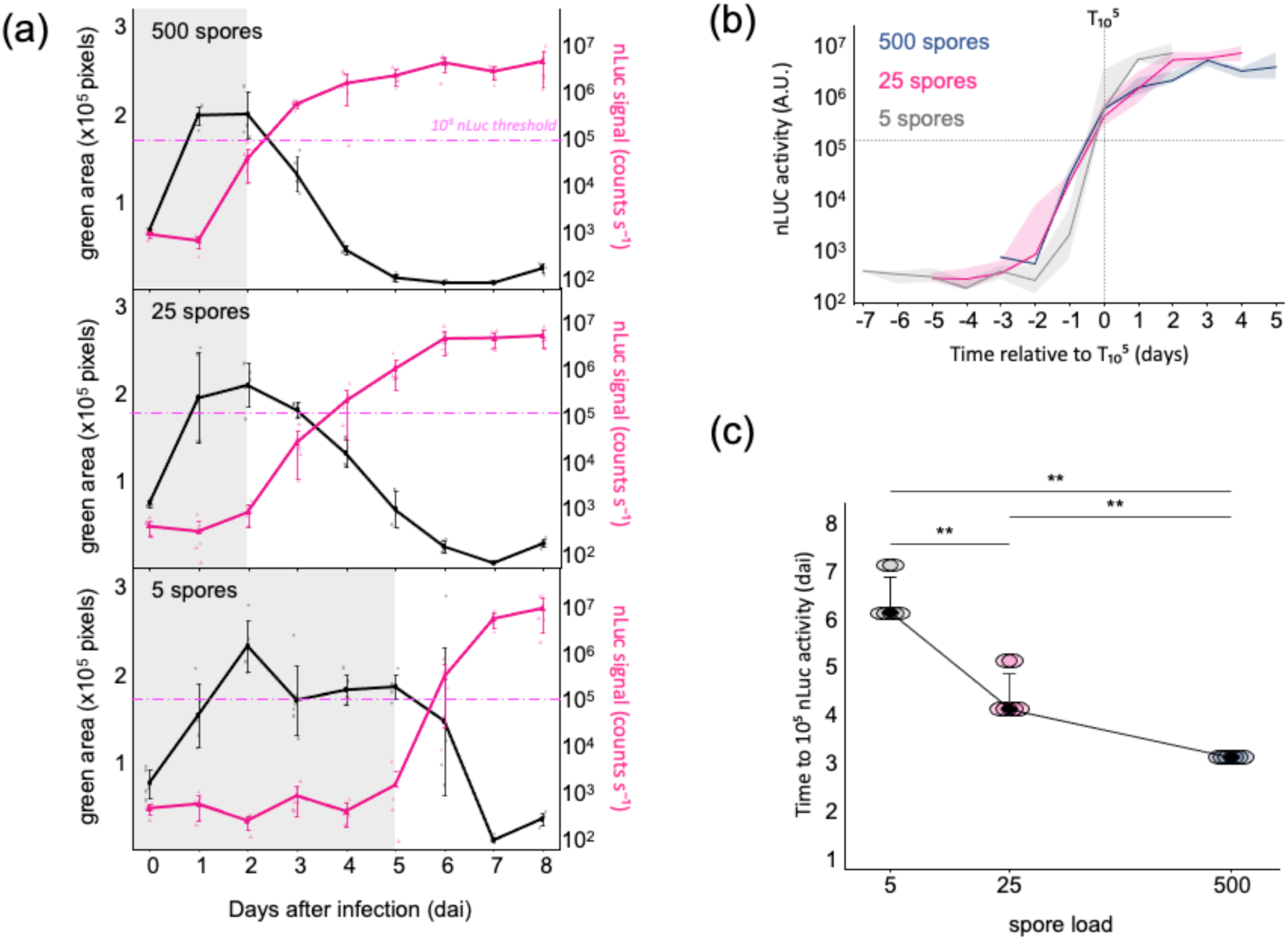
Initial spore load controls the timing, but not the subsequent kinetics. **(a)** Longitudinal monitoring of seedling green area and NanoLuc luminescence following inoculation with 500, 25 or 5 Hemera spores per inoculation site. Green area is expressed as pixel area (×10⁵ pixels), and NanoLuc luminescence is shown in arbitrary units (a.u.). Grey shading indicates the asymptomatic interval for each inoculum density from longitudinal green-area measurements. The horizontal dashed line indicates the empirically defined *10^5^*NanoLuc threshold. Increasing spore load progressively shortened the time required to reach this threshold. **(b)** Fungal expansion follows similar kinetics after 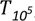. NanoLuc trajectories were aligned independently for each biological replicate to 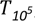, defined as the first sampling day at which NanoLuc luminescence reached *10^5^* a.u. 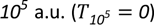. 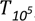 was used solely as an operational temporal reference and does not represent the biological onset of fungal growth or expansion. Solid lines represent median trajectories and shaded areas indicate the interquartile range. **(c)** Initial spore load determines the time required to reach 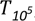. Each point represents one biological replicate and the black line connects condition medians. Differences among inoculum densities were assessed using a Kruskal–Wallis test followed by pairwise Wilcoxon tests with Holm correction. Asterisks indicate adjusted pairwise P-values: **P < 0.01.

### Early fungal progression precedes robust SA- and JA-associated defense responses

Because *Arabidopsis* seedlings continued to grow while fungal progression increased during the asymptomatic phase, we examined the contribution of the salicylic acid (SA) and jasmonic acid (JA) pathways during this early stage of infection. We used the SA-signaling mutant *npr1* and the JA-perception mutant *coi1-1* (Figure 3). Under non-infected conditions, all genotypes displayed progressive seedling growth, although *npr1* exhibited a slight developmental delay. Following inoculation, visible disease symptoms developed with broadly similar kinetics in the three genotypes. Likewise, NanoLuc bioluminescence revealed similar fungal progression in wild-type, *npr1* and *coi1-1* seedlings throughout the asymptomatic phase, indicating that impairment of either SA- or JA-dependent signaling has little effect on early fungal progression.

**Figure 3.**
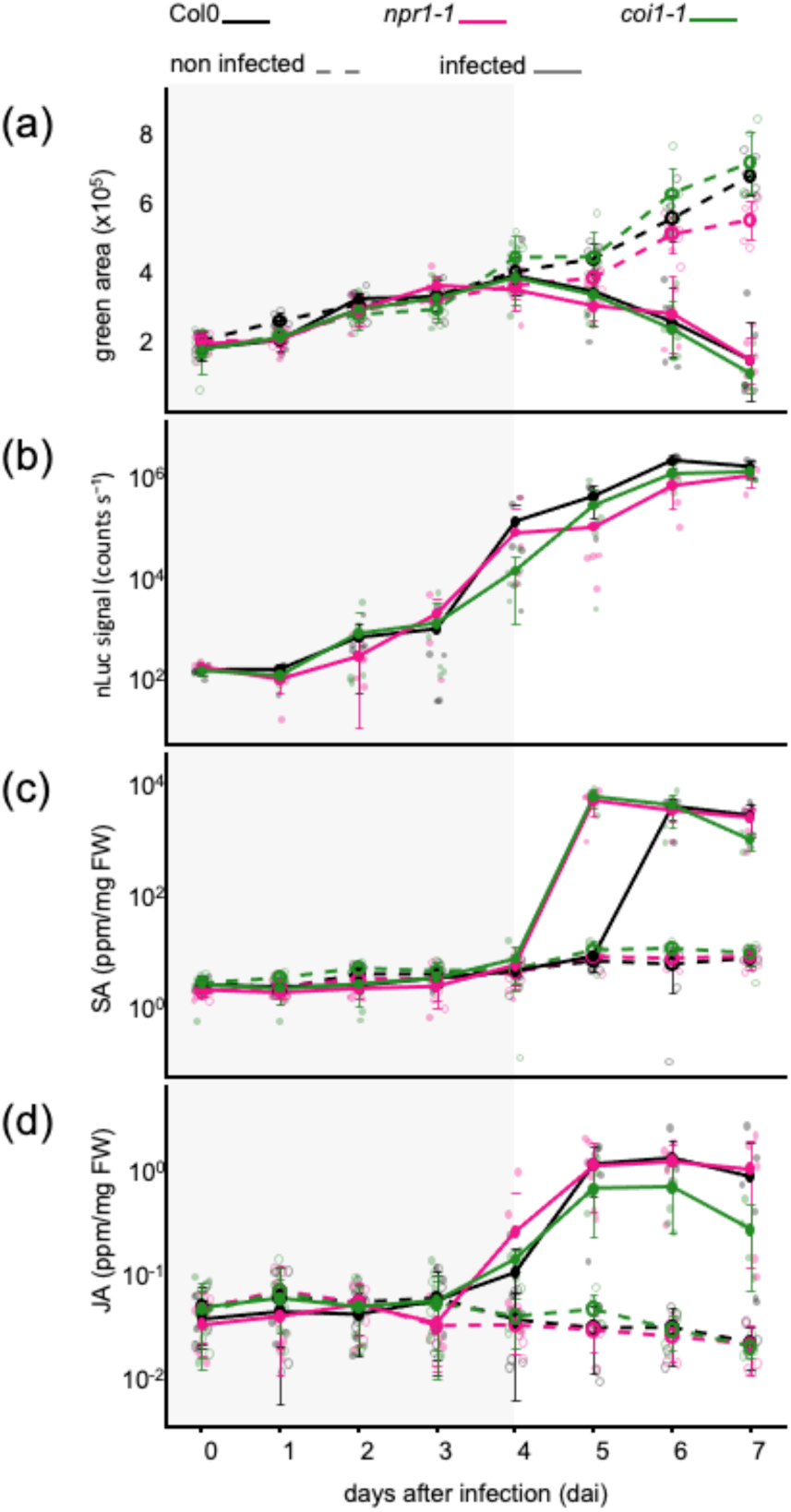
Fungal colonization precedes robust SA- and JA-associated responses in Arabidopsis seedlings. **(a)** Disease progression was monitored in Col-0, *npr1-1* and *coi1-1* seedlings by longitudinal measurement of green area. Non-infected seedlings continued to develop throughout the experiment, whereas infected seedlings showed a decline in green area from approximately 4 days after inoculation (dai). **(b)** Host-associated fungal colonization, quantified by NanoLuc luminescence, followed similar kinetics in Col-0, *npr1-1* and *coi1-1* seedlings. **(c–d)** Endogenous salicylic acid (SA) and jasmonic acid (JA) levels were quantified by LC–MS/MS throughout infection. No marked accumulation of either hormone was detected during the early phase of fungal progression. JA accumulation became apparent earlier than SA in Col-0, whereas both hormones accumulated strongly at later stages in all genotypes. Thus, substantial fungal colonization occurred before robust SA and JA accumulation.

We next quantified endogenous SA and JA levels by LC-MS throughout the infection time course (Figure 3 ; Figure S8). No significant differences in SA or JA levels were detected between infected and non-infected seedlings in any genotype up to 4 dai (Figure S9), despite the continuous increase in fungal progression during this period. JA accumulation became clearly detectable at 5 dai, reaching approximately one to two orders of magnitude above basal levels. SA accumulation occurred later but was considerably stronger, exceeding three orders of magnitude in *npr1-1* and *coi1-1* and, at 6 dai, in wild-type seedlings. Together, these results show that substantial fungal progression during the asymptomatic phase precedes robust SA and JA accumulation. Consistently, impairment of either SA or JA signaling did not markedly alter fungal progression during this early infection period.

### B. cinerea displays tissue-associated differentiation during asymptomatic colonization

To examine the cellular organization of *B. cinerea* during the prolonged asymptomatic phase, infected *Arabidopsis* seedlings were analyzed by confocal microscopy (Figure 4). Fungal morphology varied markedly with the host tissue encountered. Along the root surface, hyphae displayed a predominantly vegetative-like growth pattern, frequently winding around rhizodermal cells and occasionally forming swollen regions and discrete host-contact sites. In contrast, differentiated infection structures were frequently observed on hypocotyls and cotyledons. The hypocotyl in particular displayed diverse infection-related morphologies, including hook-shaped cells and multilobed structures, while infection cushions were also observed on aerial tissues (Figure 4). Most observed hyphae remained surface-associated, but occasional endophytic invasion events were detected in hypocotyls, with fungal hyphae passing between adjacent epidermal cells and forming multilobed structures within host tissues (Figure 4). Thus, macroscopically asymptomatic colonization encompasses markedly different local fungal morphologies, including differentiated infection structures and occasional tissue invasion.

**Figure 4.**
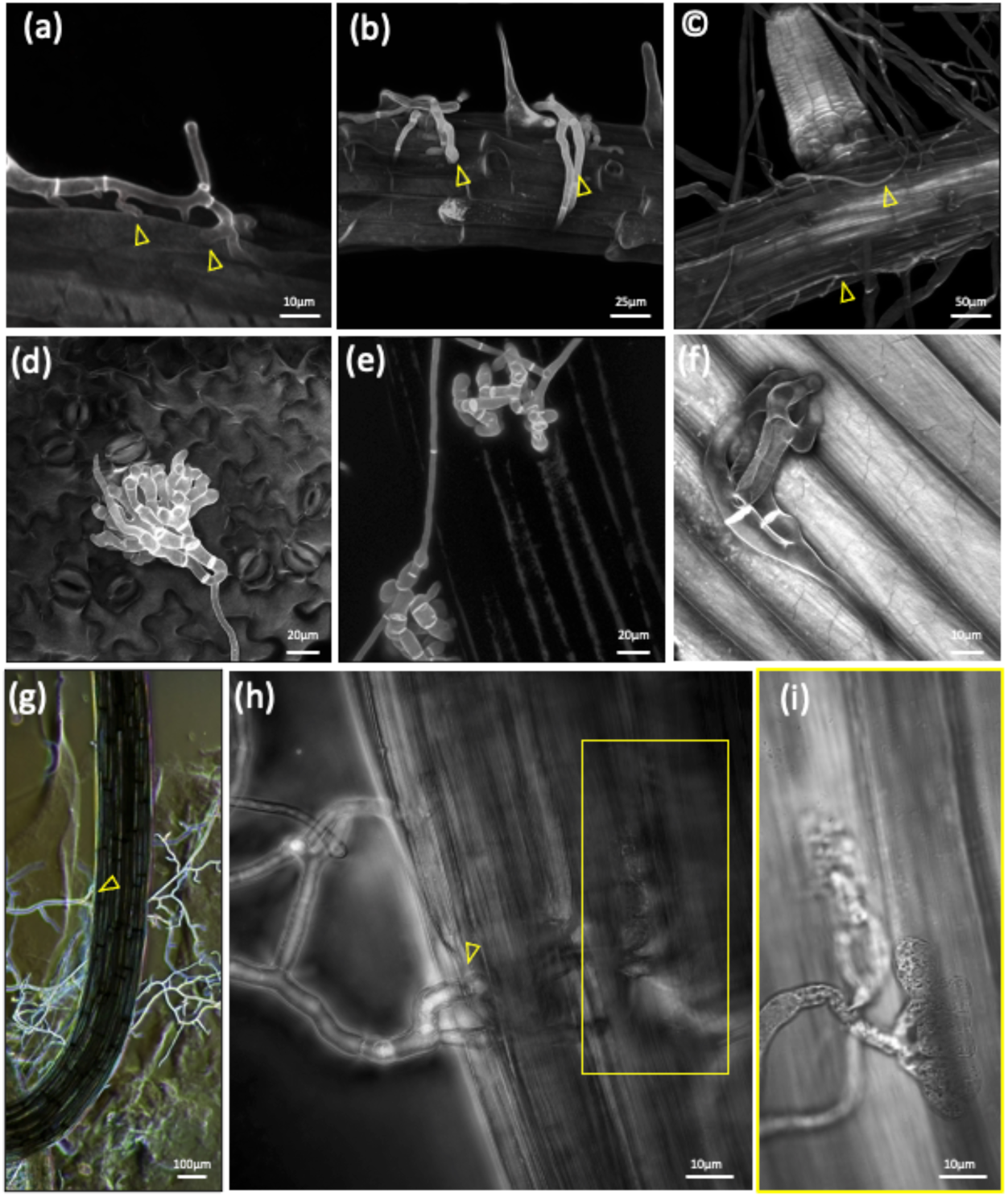
Tissue-associated infection morphologies of Botrytis during asymptomatic colonization of Arabidopsis seedlings. Arabidopsis seedlings infected with *Botrytis cinerea* were fixed directly in Petri dishes, stained with calcofluor white and imaged by confocal fluorescence microscopy following z-stack acquisition, except panels f–h, which combine bright light (f) and transmitted-light imaging to visualize fungal invasion within host tissues (g and h). **(a)** Vegetative hypha establishing discrete contact sites along the root epidermis. **(b)** Swollen vegetative hypha winding around rhizodermal cells. **(c)** Branched hyphal network surrounding the root surface. **(d)** Infection cushion developing on the cotyledon epidermis. **(e)** Differentiated multilobed infection structure associated with the hypocotyl epidermis. Pour le panel d : ajouter en commentaire que le calco colore mal la paroi des cellules de l’hypocotyl **(f)** Hook-shaped infection structure at the hypocotyl surface. **(g–i)** Endophytic invasion of the hypocotyl. A fungal hypha passes between adjacent epidermal cells and is continuous with a multilobed fungal structure located within the host tissue. Successive focal planes reveal the spatial continuity between the surface-associated hypha and the internal fungal structure. In panel h, the yellow box indicates the region shown at higher magnification in panel i.

### Spatial organization of the fungal cell wall at host contact sites

To examine fungal cell-wall organization at plant contact sites, infected seedlings were co-stained with calcofluor white and wheat germ agglutinin (WGA). Calcofluor broadly labels β-linked polysaccharides of the fungal wall, whereas WGA recognizes accessible N-acetylglucosamine residues associated with chitin. WGA strongly labelled fungal septa and discrete domains within differentiated infection structures, revealing a markedly non-uniform distribution of accessible chitin-associated epitopes (Figure 5).

**Figure 5.**
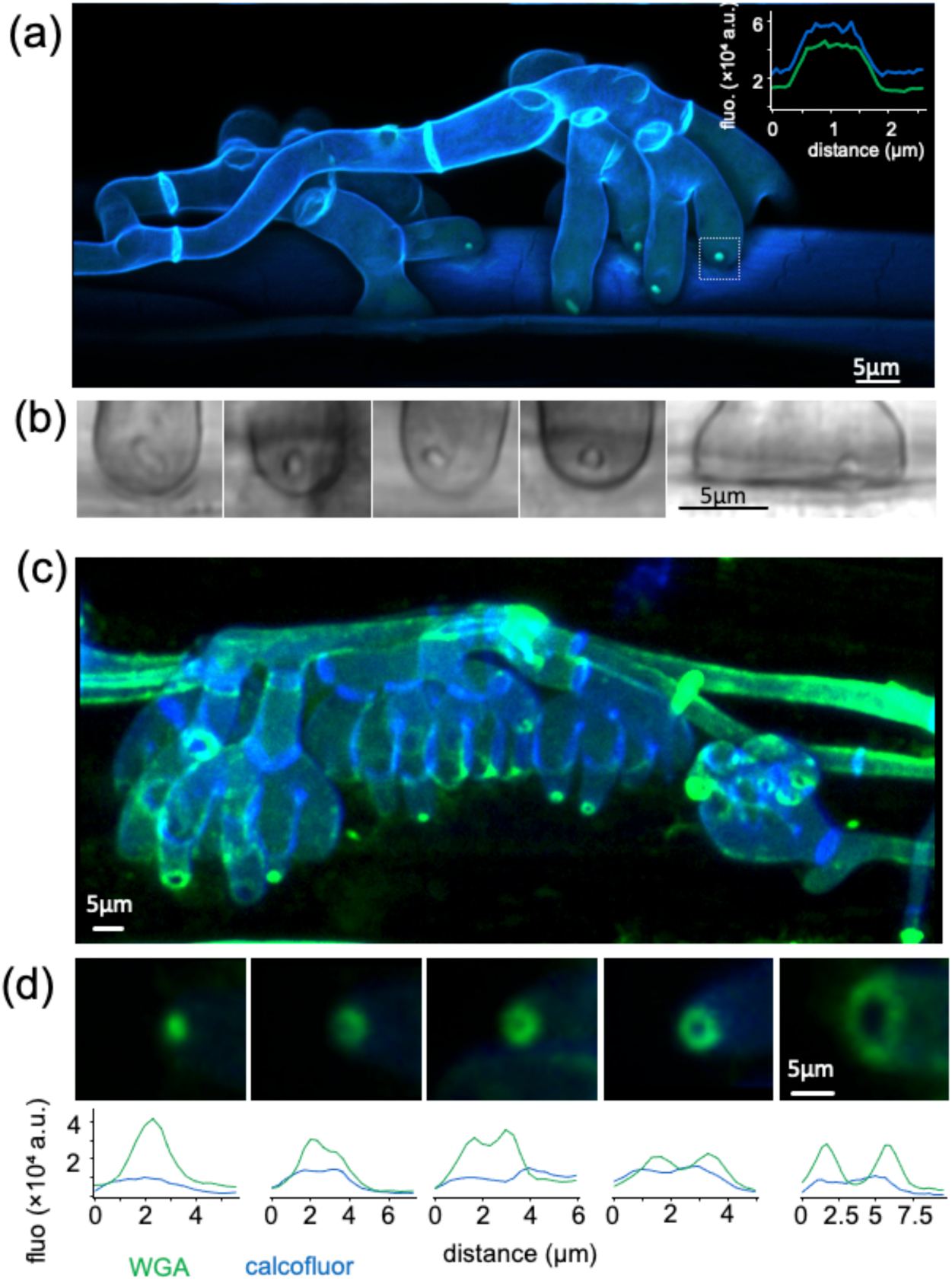
*Botrytis cinerea* infection structures display spatially heterogeneous cell-wall organization at the *Arabidopsis* interface. *Arabidopsis thaliana* seedlings infected with *B. cinerea* were fixed and co-stained with calcofluor white and fluorescent wheat germ agglutinin (WGA) before confocal z-stack imaging. Calcofluor broadly labels fungal cell walls, whereas WGA detects accessible β-1,4-linked N-acetylglucosamine residues associated with chitin. Representative fluorescence channels and merged images are shown. **(a)** Multilobed infection structures display discrete pinhead-like WGA-positive foci positioned at the extremity of individual lobes. A fluorescence intensity line profile across a representative terminal domain compares the spatial distribution of WGA and calcofluor signals. **(b)** Transmitted-light microscopy of representative lobe extremities reveals morphologically distinct terminal domains corresponding to the WGA-enriched structures observed by fluorescence microscopy. **(c)** Correlative observation of the same infected hypocotyl (3 dai) after fixation. The seedling was first imaged by stereomicroscopy (figure 2g,h) and subsequently fixed, co-stained with calcofluor white and fluorescent wheat germ agglutinin (WGA), and examined by fluorescence microscopy. Punctate and ring-shaped WGA-positive domains are observed along the fungal surface at host contact sites. **(d)** Higher-magnification views of representative punctate and ring-shaped domains, together with corresponding fluorescence intensity line profiles of WGA and calcofluor signals, illustrate their distinct spatial distributions at fungal–plant contact sites. Together, these observations reveal pronounced spatial heterogeneity in the organization and accessibility of chitin-associated cell-wall domains within differentiated *B. cinerea* infection structures.

At fungus–plant contact sites, WGA-positive domains displayed several recurrent spatial patterns. Multilobed infection structures frequently contained small pinhead-like WGA-positive foci at the extremity of individual lobes (Figure 5a). These terminal domains were also visible by transmitted-light microscopy, indicating that WGA enrichment coincided with morphologically differentiated fungal structures at the host surface (Figure 5b). Other infection structures displayed punctate or ring-shaped WGA-positive domains closely associated with the host-contacting fungal surface (Figure 5 c.d). Fluorescence intensity profiles across representative domains showed sharp local WGA enrichments that were not mirrored by the broader calcofluor signal (Figure 5 a.d).

Although the relationship between these different patterns remains unresolved, their recurrent localization at fungal contact sites reveals pronounced spatial heterogeneity in the organization and accessibility of chitin-associated cell-wall domains at the *B. cinerea*–host interface.

### Membrane and cytoskeletal remodeling at the *Botrytis–Arabidopsis* interface

The close association between differentiated fungal structures and host cells raised the question of whether the *Arabidopsis* cortical architecture is locally remodeled at these contact sites. To visualize fungus–plant interfaces by live-cell confocal microscopy, we established a slide– coverslip pathotest compatible with high-resolution imaging of *Botrytis–Arabidopsis* interaction sites (Figure 6 a).

**Figure 6.**
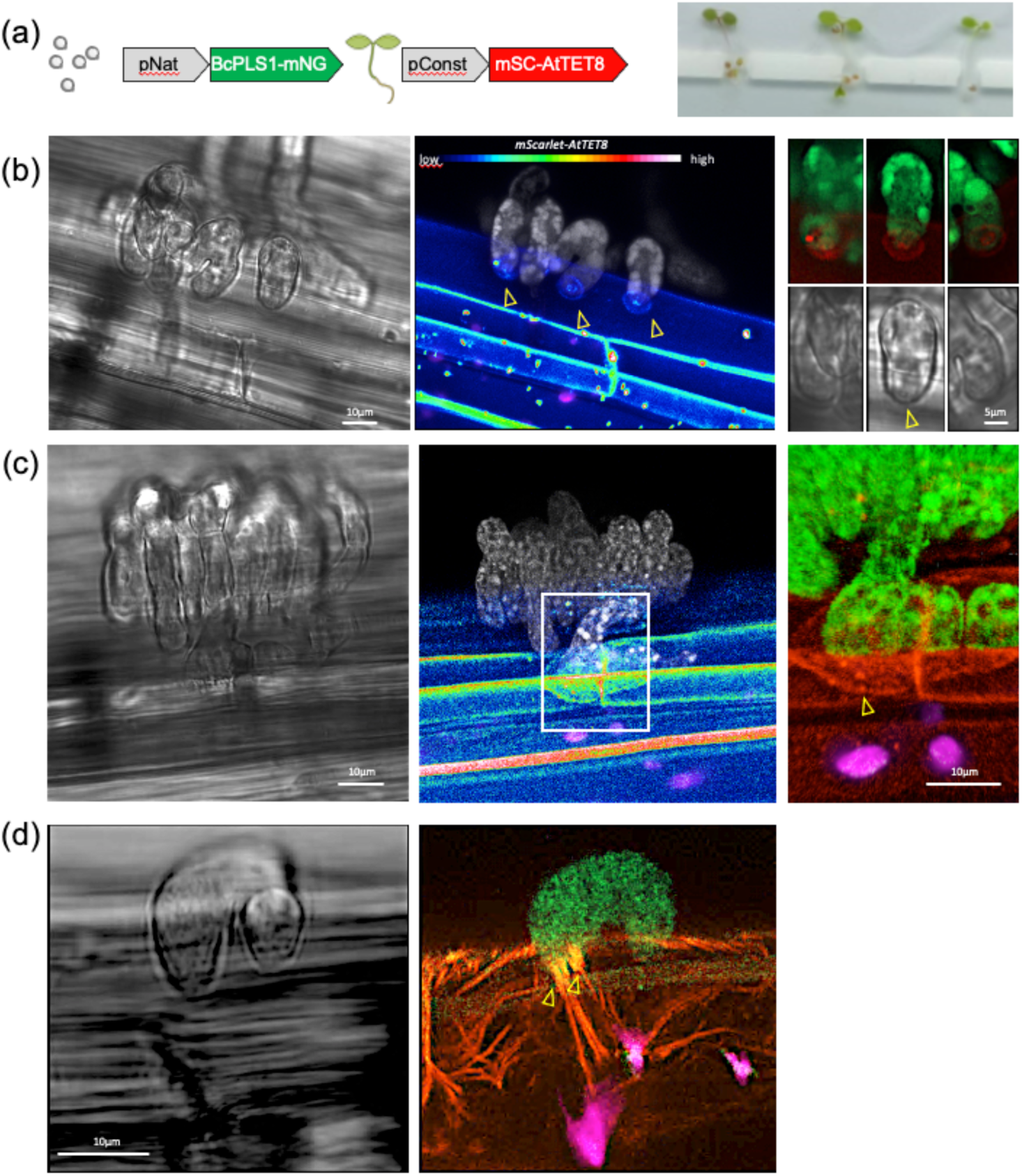
*Arabidopsis* cells undergo localized membrane and actin polarization at the *Botrytis* interface. Localized TET8 accumulation at the plasma membrane and cortical actin reorganization are associated with differentiated fungal contact sites. **(a)** Live-cell imaging setup combining *Arabidopsis thaliana* hypocotyl cells constitutively expressing mScarlet-TET8 with a *Botrytis cinerea* strain expressing the tetraspanin reporter PLS1-mNeonGreen. Infected seedlings were mounted between a slide and coverslip to allow confocal visualization of fungus–host interfaces during the first two to three days after inoculation. **(b)** Representative differentiated fungal structure in contact with the hypocotyl epidermis. Transmitted-light imaging shows the fungal morphology at the host surface. PLS1-mNeonGreen identifies the differentiated fungal structure, whereas mScarlet-TET8 accumulates locally at the host plasma membrane facing individual fungal lobes. Higher-magnification views highlight localized TET8-enriched membrane domains associated with fungal contact sites. A pin-like fungal structure is indicated by an arrow. mScarlet-TET8 intensity is displayed using a rainbow lookup table (LUT), with PLS1-mNeonGreen shown in white. **(c)** Independent example of an infection cushion contacting hypocotyl epidermal cells. mScarlet-TET8 signal increases locally at the plasma membrane directly facing the fungal structure, with discrete TET8-enriched foci visible at the contact site (arrow). **(d)** *Arabidopsis* LifeAct reporter showing pronounced reorganization of the cortical actin network at a differentiated fungal contact site. Actin bundles converge toward the fungal structure, with the arrow indicating a prominent site of actin convergence at the fungus–host interface. All images were acquired from hypocotyl tissues. Scale bars are indicated in the corresponding panels.

To distinguish fungal infection structures from host membrane responses, we combined an *Arabidopsis* line constitutively expressing mScarlet-TET8 with a *B. cinerea* strain expressing the tetraspanin reporter PLS1-mNeonGreen, which preferentially labels differentiated infection structures (Figure 6 a; Figure S10, S11). In non-infected seedlings, TET8 was predominantly associated with the plasma membrane. During infection, TET8 frequently accumulated in discrete cortical domains directly facing differentiated fungal structures (Figure 6 b.c; Figure S11). These enrichments ranged from small localized domains to larger ring-like assemblies outlining fungal contact sites. In several interaction events, host TET8- enriched domains were closely apposed to PLS1-positive fungal structures, revealing a highly localized organization on both sides of the interface.

Because cortical actin is central to membrane trafficking and polarized cellular responses to external cues, we next asked whether TET8 polarization was accompanied by cytoskeletal reorganization. Imaging of *Arabidopsis* LifeAct reporter lines revealed pronounced local remodeling of the cortical actin network beneath fungal contact sites. Actin bundles repeatedly converged toward differentiated infection structures and occasionally formed dense cortical accumulations immediately beneath fungal cells (Figure 6 d; Figure S11). Similar actin polarization was observed in several independent interaction events while surrounding host cells remained morphologically intact, showing that localized cytoskeletal reorganization can occur in host cells without overt signs of cellular collapse at the time of imaging.

Collectively, TET8 and LifeAct reporters revealed localized remodeling of the host plasma membrane and cortical cytoskeleton at fungal contact sites. TET8 accumulated in discrete membrane domains, whereas cortical actin repeatedly polarized toward differentiated infection structures. These responses were observed in host cells displaying an apparently continuous plasma membrane together with preserved chloroplast morphology and chlorophyll autofluorescence, consistent with maintained cellular integrity at the time of observation.

Together with the spatially patterned WGA-positive wall domains observed on the fungal side, these findings reveal highly organized fungus–plant interfaces associated with differentiated infection structures. Local TET8 enrichment and actin polarization demonstrate active remodeling of the host cortical architecture, while the fungal contact surface displays a spatially heterogeneous cell-wall organization. Thus, differentiated *B. cinerea* structures are associated with strongly polarized interactions with morphologically intact host cells at the cellular scale.

## Discussion

### Active fungal colonization precedes widespread host tissue collapse

Our results reveal a prolonged infection window during which substantial host-associated fungal colonization precedes widespread tissue collapse. Colonization increased strongly while *Arabidopsis* seedlings continued to expand their green tissues, showing that the absence of macroscopic symptoms did not reflect fungal quiescence. At the scale of the whole seedling, this interval therefore defines a macroscopically asymptomatic phase in which substantial fungal colonization remains compatible with continued host development.

This temporal separation also extended to host hormonal responses. Strong whole-seedling accumulation of JA and SA occurred mainly at later stages, whereas fungal colonization had already increased substantially. Several non-exclusive fungal mechanisms could delay or spatially restrict immune outputs during early colonization. Across phytopathogenic fungi, immune recognition can be reduced by masking or modifying immunogenic cell-wall components, while secreted effectors interfere with host signalling. In *B. cinerea*, BcLysM1 limits chitin-triggered immunity, and fungal sRNAs can hijack host AGO1 to silence immunity-related genes (Weiberg *et al.,* 2013; Crumière *et al.,* 2025). Modification of hormone homeostasis provides another possible mechanism, as fungal catabolism of SA has been demonstrated in *Ustilago maydis* and *Sclerotinia sclerotiorum* (Rabe *et al*., 2013; Penn & Daniel, 2013).

The absence of robust early SA and JA accumulation at the whole-seedling scale does not, however, imply that the host fails to perceive the fungus. High-resolution imaging revealed pronounced local plasma-membrane and cortical-actin reorganization at individual fungus– plant contact sites. Early responses may therefore remain spatially restricted and become diluted in measurements performed on entire seedlings. Because hormone profiling and interface imaging were performed using complementary plate and slide–coverslip assays, respectively, their precise temporal equivalence cannot be established, but together they highlight an important distinction between global hormonal outputs and localized cellular responses.

Fungal colonization was also broadly similar in wild-type, *npr1-1* and *coi1-1* seedlings during the early infection period, suggesting that NPR1-dependent SA signalling and COI1-dependent JA perception do not impose a major net restriction on colonization under our conditions. This does not imply pathway inactivity. Co-transcriptomic analyses have shown that *npr1-1* and *coi1-1* backgrounds extensively reshape early *B. cinerea* transcriptional programmes, including networks associated with fungal growth, membrane and vesicle functions, secretion and virulence (Zhang *et al.,* 2019). Those experiments were performed on detached leaves of five-week-old plants, emphasizing the different developmental context of our seedling system.

Asymptomatic *Botrytis* interactions may therefore encompass different immune and hormonal states rather than a single weakly responsive condition. Selected non-pathogenic interactions in tomato can induce resistance that depends primarily on SA and ethylene signalling (Anand *et al*., 2026), while immune competence itself changes during plant development. In *Arabidopsis*, FLS2-mediated immunity increases during early seedling development, and susceptibility to *B. cinerea* varies with leaf developmental stage in tomato (Zou *et al.,* 2018; Lindner *et al*., 2025). The macroscopically asymptomatic phase observed here is therefore neither immunologically silent nor readily explained by developmental incompetence alone.

### Asymptomatic colonization encompasses context-dependent infection-related fungal states

A striking feature of this phase was the diversity of fungal morphologies associated with different host tissues. Vegetative-like hyphae, hook-shaped and multilobed structures, infection cushions and occasional invasive structures were observed before widespread visible disease. Asymptomatic colonization is therefore not equivalent to a uniform pre-infectious state; vegetative-like and differentiated infection-related morphologies coexist during this phase. Fungal morphology also differed among host tissues. Roots were predominantly colonized by surface-associated hyphae, whereas hypocotyls and cotyledons displayed a broader range of differentiated infection structures. Because these observations are qualitative, they do not demonstrate tissue-specific developmental programmes. They are nevertheless consistent with fungal plasticity across distinct host microenvironments, without implying that any particular morphology is specifically adapted to a given tissue.

The extensive association with roots is noteworthy in light of the broader ecology of *B. cinerea*. Symptomless systemic colonization of lettuce has been documented from roots through stems and leaves as plants develop (Sowley *et al*., 2010), and fungal inoculum can persist in soil-associated plant debris and long-lived sclerotia (Araújo *et al*., 2005; Leyronas *et al.,* 2015). This does not imply that root-associated growth in our system is saprotrophic, but it emphasizes that roots constitute one of several nutritionally and physiologically distinct environments potentially encountered by the fungus. Host development can further modify the outcome of colonization: in *Primula*, latent infections established in young plants can remain symptomless for extended periods, with subsequent symptom expression strongly influenced by plant development (Barnes & Shaw, 2002). Infection outcome may therefore remain contingent on tissue identity and host developmental state well after initial colonization.

The functional significance of the fungal morphologies observed here remains unresolved. Different structures may represent sequential stages of a common developmental programme, alternative responses to local host conditions or distinct fungal functional states. Work in other appressorium-forming fungi supports a distinction between morphological differentiation and acquisition of full invasive competence. In *Magnaporthe oryzae*, hierarchical regulation downstream of Pmk1 separates early appressorium differentiation from later programmes controlling repolarization, cytoskeletal organization, secretion and invasive growth (Osés-Ruiz *et al*., 2021). Host-derived metabolites can directly redirect fungal development. Triterpenoid saponins alter the balance between epiphytic growth, unicellular appressorium formation and subsequent infection-cushion development in *B. cinerea* (Escaray *et al.,* 2024). Developing seedlings are particularly relevant in this respect because roots, hypocotyls and cotyledons differ in architecture, physiology and specialized metabolism. *Arabidopsis* roots themselves produce a spatially organized repertoire of specialized triterpenes, some of which occur at the root surface and are released by seedlings (Huang *et al*., 2019). Their influence on *B. cinerea* remains unknown, but such chemical heterogeneity provides a plausible mechanism through which host context could influence fungal differentiation.

For a broad-host-range pathogen, this developmental plasticity could facilitate colonization across heterogeneous host environments. Our observations also suggest that morphology alone may not predict the functional state of an infection structure. Whether the forms observed here represent sequential maturation stages, distinct fungal states or a reversible continuum remains unresolved.

### Differentiated fungal structures are associated with spatially organized plant–fungus interfaces

Differentiated fungal structures frequently formed close contacts with plant cells retaining viability-associated features. On the fungal side, WGA staining revealed discrete pinhead-like, punctate and ring-shaped domains at contact sites. Chitin accessibility can be reduced through complementary mechanisms. α-1,3-glucan masks surface chitin in *Magnaporthe oryzae*, *Cochliobolus miyabeanus* and *Rhizoctonia solani* (Fujikawa *et al*., 2012), while fungal LysM effectors can bind chitin, protect hyphae from chitinases and limit chitin-triggered immunity (de Jonge *et al*., 2010; Mentlak *et al*., 2012). In *B. cinerea*, these functions are carried by BcLysM1 (Crumière *et al*., 2025). Because WGA reports probe-accessible GlcNAc-containing material rather than total fungal chitin, local WGA enrichment should not be interpreted as a direct measure of increased chitin abundance, nor should WGA-poor regions be interpreted as chitin depletion. Instead, these patterns reveal spatial heterogeneity in cell-wall organization and accessibility, the molecular basis of which remains unresolved.

The plant compartment was also strongly polarized. TET8-containing plasma-membrane domains accumulated as discrete foci or larger assemblies facing differentiated fungal cells, while cortical actin locally converged toward fungal contact sites. TET8 recruitment to B. cinerea infection sites was previously reported in the context of multivesicular-body trafficking and extracellular-vesicle-mediated transfer of host sRNAs (Cai *et al*., 2018). A similarly focal redistribution of TET8 has been observed during powdery mildew infection, where TET8-GFP accumulates at successful penetration (Rubiato *et al*., 2022). Our imaging extends these observations by resolving sharply localized TET8 domains at individual B. cinerea–plant interfaces. TET8-like tetraspanin polarization is not restricted to pathogenic interactions. In Phaseolus vulgaris, PvTET8-1 accumulates at the apical membrane of growing root hairs and at sites of redirected polar growth, and its manipulation affects both rhizobial nodulation and arbuscular-mycorrhizal colonization (Parra-Aguilar *et al.,* 2023). Together, these observations suggest that TET8-like tetraspanins participate more broadly in the organization of specialized membrane domains at intimate plant–microorganism interfaces, rather than functioning exclusively as markers of defensive EV secretion. Together, these observations suggest that TET8 recruitment reflects a broader organization of specialized plasma-membrane domains at fungal contact sites, beyond its established association with extracellular vesicle trafficking.

Across our complementary imaging approaches, both fungal and host compartments displayed pronounced spatial organization at contact sites: fungal wall accessibility was patterned, whereas TET8-containing membrane domains and the cortical actin network underwent focal reorganization toward differentiated fungal structures. These observations support the formation of a polarized cellular interface and are compatible with localized secretion, endocytosis or other forms of membrane trafficking.

What sustains *B. cinerea* colonization during the macroscopically asymptomatic phase?

Low inoculum density markedly prolonged the pre-expansion interval, whereas post-threshold kinetics were comparatively similar, particularly between the 25 and 500 spore treatments. Low spore loads may therefore expose a phase that is temporally compressed and difficult to resolve at higher inoculum densities. Importantly, the subsequent rapid increase in fungal colonization occurred while the host remained macroscopically asymptomatic.Under these experimental conditions, fungal growth does not necessarily require direct exploitation of host tissues. *B. cinerea* was able to grow slowly on the sucrose- free MSk medium in the absence of seedlings, showing that the experimental substrate itself can support a limited degree of fungal growth. During infection, additional resources could derive from compounds released by the plant, including exudates, which can represent substantial carbon pools for plant-associated microorganisms (Canarini et al., 2019), as well as from localized tissue damage or nutrient redistribution through the connected fungal network. The relative contribution of these sources cannot be resolved from the present experiments. Fungal colonization may be sustained by a combination of extrahost and host-associated resources, whose relative importance may change as infection progresses. A contribution from nutrients available outside living plant cells would also be consistent with the ecological versatility of *B. cinerea*, which can alternate between pathogenic growth and saprotrophic exploitation of non-living plant material. Determining the carbon sources supporting fungal growth during this phase will be essential for defining its trophic basis.

### Concluding remarks and perspectives

This study establishes a multiscale framework for resolving B. cinerea colonization during a macroscopically asymptomatic phase that is difficult to capture in conventional detached-leaf assays. Intact developing seedlings, low spore loads and sucrose-free conditions allowed fungal colonization and host development to be followed longitudinally, while complementary microscopy approaches provided access to individual fungus–plant interaction sites. Combined with the Hemera reporter, this system also provides a scalable platform for testing how fungal genotype, plant genotype, developmental stage and environmental conditions influence colonization and disease progression.

Together, our results identify a macroscopically asymptomatic phase in which B. cinerea undergoes substantial colonization and infection-related differentiation while host development continues. At the cellular scale, differentiated fungal structures are associated with highly organized plant–fungus interfaces, characterized by spatially patterned fungal cell-wall domains and focal reorganization of the host plasma membrane and cortical actin network. These observations raise the question of the molecular dialogue operating across these interfaces and of whether the distinct fungal morphologies observed here correspond to distinct functional cell states.

Addressing these questions will require moving beyond morphology toward cell-resolved molecular approaches. Single-cell or single-nucleus transcriptomics (Zhu et al., 2023), ideally combined with spatially resolved imaging and molecular reporters, should help determine whether vegetative hyphae, multilobed structures, infection cushions and invasive cells correspond to distinct transcriptional programmes during asymptomatic colonization. Applying comparable approaches to the facing plant cells would further reveal whether localized membrane and cytoskeletal polarization is associated with specific host transcriptional states. Integrating the molecular states of both partners at individual infection sites should ultimately clarify how local fungus–plant interfaces are established, maintained and reprogrammed as asymptomatic colonization progresses toward destructive disease.

## Material and methods

### Biological material

*B. cinerea* strains are detailed in Table 1. *Agrobacterium tumefaciens* LBA1126 was used for *B. cinerea* transformation and GV3101 (pSOUP) for *A. thaliana* transformation. *A. thaliana* lines are summarized in Table 2; *npr1-1* and *coi1-1* were provided by M. Fagard (IJPB, INRAE, Centre Ile-de-France Versailles-Saclay).

### Plasmids

pFX217 (PoliC::nLuc, *nptII*; flanked by the *BcniaD*-*Bcsrp68* intergenic locus) was used for *B. cinerea* transformation and pFX131 (p35S::AtTET8-mScarlet, pGreenII backbone) for *A. thaliana* transformation (Table 3, Fig. SX).

### *Botrytis cinerea* spore preparation

Strains were grown on malt sporulation medium (table 6) supplemented with the selection antibiotic (neomycin 150 µg/mL or hygromycin 70 µg/mL) under dark conditions (2 days, 21 °C), then exposed to near-UV light (365 nm, 9 days, 21 °C) to induce sporulation. Spores were harvested in 15% glycerol, 40 µm filtered to remove mycelial debris, counted on Thoma cells, adjusted to 10^7 spores/mL and stored at −80 °C as single-use aliquots. Post-freeze germination was checked for each batch on MS agar (1-2 days; acceptance threshold 95%).

### *Agrobacterium tumefaciens*-mediated transformation (ATMT)

Plasmids were introduced into *A. tumefaciens* LBA1126 competent cells by heat shock (5 min, 37 °C), transformants were selected on Lysogeny Broth (LB) low-salt medium containing kanamycin (50 µg/mL) and spectinomycin (250 µg/mL) (2-3 days, 28 °C), positive colonies were confirmed by PCR and stocked in 15% glycerol at −80 °C (Barbara Schrammeijer *et al*., 2003). Transformation of B05.10 was performed as previously described (Rolland *et al*., 2003). *A. tumefaciens* was grown in induction medium (IM, Table 6) to OD600 0.4-0.5, mixed (1:1, v/v) with a spore suspension (10^7 spores/mL) in liquid IM (5 min, RT), spread on cellophane sheets (#1650963 Bio-Rad) overlaying solid IM (150 µL/plate) and incubated (48 h, 21 °C). Transformants were selected on TNK-YE glucose medium with neomycin (500 µg/mL) and cefotaxime (200 µM) (3- 7 days, 21 °C), submitted to two rounds of selection on MEA with the same antibiotics (4-5 days each) and maintained on Potato Dextrose Agar (PDA, Millipore) with neomycin (150 µg/mL). Monospore isolation was performed on malt sporulation medium (3% agar) (16 isolates per strain; 5-7 days, 20 °C, dark). Fungal genomic DNA was extracted in 20 mM NaOH (20 min, 98 °C) and recombination junctions and ectopic insertions were checked by PCR (Table 4).

### *Arabidopsis thaliana* seeds sterilization and plate pathotest preparation

*A. thaliana* seeds sterilization was performed using a sterilization solution (bleach, absolute ethanol, Tween 20; Table 6) for 5 min, followed by two washes in 70% ethanol and one wash in absolute ethanol (2 min each). Seeds were dried and spread onto 100 µm nylon membranes (1.5 x 1.5 cm; ∼55 seeds per membrane) using a 1 mm mosquito-net sowing grid. Nylon membranes were then transferred onto 55 mm Murashige and Skoog plates (MSk, Duchefa Biochemie #M0231.005; 8 mL, 0.9% plant agar, pH 5.7). After stratification (3 days, 4 °C, dark), germination induction (6 h, 21 °C, light), skotomorphogenesis (3 days, 21°C, dark) and photomorphogenesis (2 days, 21°C, light), a single 10 µL droplet of *B. cinerea* spore suspension (2.5 x 10^3 spores/mL, ∼25 spores) was deposited in the center of each nylon membrane under constant magnetic stirring of the inoculum, and plates were kept in open crystal boxes in a phytotron (21 °C, 40 % RH, 16 h/8 h, 70 µE/m2/s). Delivered dose was controlled by counting 20 independent 10 µL aliquots per session. Pathotests were monitored over 7 days after infection (dai) with six biological replicates per condition. Spore load experiments were performed identically with suspensions delivering 5, 25 or 500 spores per plate. At each sampling time point, only seedling material was harvested; the surrounding agar and fungal mycelium extending beyond the seedlings were excluded. Thus, qPCR and NanoLuc measurements specifically reflect plant-associated fungal material rather than total fungal growth within the plate.

### Colorimetric analysis

Plates were photographed daily under the microbiological safety cabinet using a smartphone fixed above a black and white Plexiglas support, illuminated by a LED ring light at constant mode and intensity, and triggered by a Bluetooth remote shutter (lid removed, no zoom, no filter). Images were processed with GreenDayz (Tguafaiti et al., 2026), a Python application: images were cropped to the Petri dish region and corrected for acquisition-related colour variation using the white reference area surrounding the dish. Green tissue was segmented in the HSV colour space (hue interval, minimum saturation, minimum brightness), with identical thresholds within an experiment and visual validation of mask overlays. Green area is reported as the projected number of green-mask pixels and is interpreted as an operational phenotype, not as biomass or chlorophyll content.

### qPCR analysis

Seedlings were harvested in 2 mL tubes containing one stainless-steel bead, flash-frozen in liquid nitrogen and stored at −80 °C. Samples were lyophilised (24 h), weighed and ground using a TissueLyser II (QIAGEN; 5 min, 30 s-1). Genomic DNA was purified (ChargeSwitch gDNA Plant Kit, Invitrogen #CS18000), quantified by Qubit (dsDNA BR or HS assays, Invitrogen #Q32850 and #2409104) and stored at −20 °C. Absolute assays targeted the nuclear single-copy locus *BcPDA1* (Bcin07g01890) and the mitochondrial locus *BcCYT* (BC1G_20024); primers were designed with NCBI Primer-BLAST and validated as shown in Figure S3. Amplicon standards were gel-purified (QIAquick Gel Extraction Kit, QIAGEN #28704) and quantified by Qubit. qPCR was performed in 96-well plates with PowerUp SYBR Green Master Mix (Applied Biosystems) on an Applied Biosystems 7900HT (experiment 1) and on Biorad *CFX Opus* (experiment 2), using ten-fold serial dilutions of each amplicon (10^-2 to 10^-7 ng/µL), triplicate standards and samples, and no-template controls. Standard curves were obtained by linear regression of Cp against log10 copy number (Table 5).

### Bioluminescence analysis

Seedlings were harvested daily from pathotest plates over 7 dai, flash-frozen in liquid nitrogen and stored at −80 °C. Seedlings were lyophilised for 1 day and ground using a tissue lyser (one 2.8 mm ceramic bead per tube) for 5 min at frequency 30/s. The powder was resuspended (1:20, w/v) in the lysis buffer (Table 6), before vortexing (1 min; RT) and centrifuged (13,000 x g, 10 min, 4 °C). 50 µL of each sample (diluted 1:10 in the lysis buffer) were transferred into a microplate well. Five microliters of detection solution (Nano-Glo HiBiT, Promega #N3030) consisting of HiBiT substrate (1:30) in a lysis buffer was added per well before measurement using a Spark microplate reader (Tecan) (Hall *et al*., 2012). Activity was normalized to the background measured in non-infected seedlings.

### Hormone extraction and quantification

The method has been adapted from Pan *et al*. (2010). Freeze-dried seedlings were weighed in pierced 2 mL tubes and ground in a TissueLyser (30 Hz, 5 min, 2.8 mm ceramic beads, adapter blocks pre-cooled at −80 °C). Hormones were extracted in 100 µL of 2-propanol/H₂O/HCl 12 N (2:1:0.002, v/v/v), without the dichloromethane partition and evaporation steps. Samples were rotated (6 rpm, 30 min, 4 °C), centrifuged (16,000 × g, 5 min, 4 °C), and 50 µL of supernatant was transferred to vials. Salicylic acid (SA) and Jasmonic acid (JA) were quantified by UPLC-ESI-MS/MS on a Waters Xevo TQ-S micro fitted with an Acquity BEH C18 column (50 × 2.1 mm, 1.7 µm, 40 °C) at 0.8 mL min⁻¹, with H₂O and acetonitrile, both containing 0.1 % formic acid, as mobile phases A and B (linear gradient from 95 % to 2.5 % A in 1 min, held 0.2 min; run time 1.5 min; injection volume 5 µL). Detection was in negative electrospray mode (capillary 3.0 kV; desolvation 600 °C and 1,200 L h⁻¹; cone gas 120 L h⁻¹), monitoring the transitions *m/z* 137.0 → 92.8 for SA and *m/z* 209.0 → 59.0 for JA (cone 10 V, collision energy 15 V, dwell time 0.164 s). Quantification was based on an external calibration curve (0.5, 1, 2.5, 5, 10, 25 and 50 ppb) run before and after each batch of 48 samples, with solvent blanks interleaved; limits of quantification were 0.1 ppb for SA and 0.05 ppb for JA. Concentrations were normalised to sample dry mass (Figure S8).

### Slide-coverslip pathotest

A custom imaging chamber was prepared on a 20 × 76 mm microscope slide using two layers of Parafilm M as a frame and spacers, resulting in an approximate chamber depth of 260 µm. Slides were sterilized in 70% ethanol for 10 min and dried under sterile conditions. The lower part of the chamber was filled with 100 µL of viscous MSk medium containing 0.1% plant agar (1 g L⁻¹; pH 5.7). Elongated *Arabidopsis thaliana* seedlings at the cotyledon stage were transferred individually onto the slide, and each hypocotyl was inoculated with a 3 µL drop of *B. cinerea* spore suspension containing approximately 5 conidia. A 5 µL drop of 0.1% agar MSk medium was added at the upper edge of the chamber to limit dehydration, and the preparation was closed with a sterile 24 × 50 mm coverslip. Mounted slides were incubated vertically in a sterile glass slide-staining tank maintained under humid conditions with sterile water and sealed with Parafilm. Fungus–plant interaction sites were examined by confocal microscopy between 2 and 4 days after inoculation.

### Plant tissue fixation and fluorescent labeling

Buffers were filtered at 0.2 µm before use (Table 6). Infected seedlings were fixed directly in Petri dishes in PME fixation solution under vacuum (2 x 5 min), treated with PBS-borohydride (5 min, vacuum) and washed three times in PBS-sorbitol (5 min each, vacuum). For labeling, samples were rehydrated through a glycerol series (PBS-glycerol 30%, 15%, then PBS 1X; 30 min each, 4 °C) and incubated with WGA-Alexa Fluor 488 (5 µg/mL) and propidium iodide (10 µg/mL) in PBS 1X (30 min RT or overnight 4 °C, gentle agitation), then washed three times in PBS 1X (10 min each). Calcofluor white (10 µg/mL) was added before observation. Specimens were mounted between slides and coverslips using adhesive chambers.

### Confocal microscopy

Confocal acquisitions were done using a Zeiss LSM 810 confocal microscope (63x oil-immersion objective) and Zen software (Zeiss - Zen blue light). Fluorescence signals were detected using the excitation (lex) and emission (lem) as described in Table 7. Laser power, gain, pinhole, detector settings, z-step and resolution were adjusted when necessary to optimize image quality, while ensuring comparability.

---

**Figure S1.**
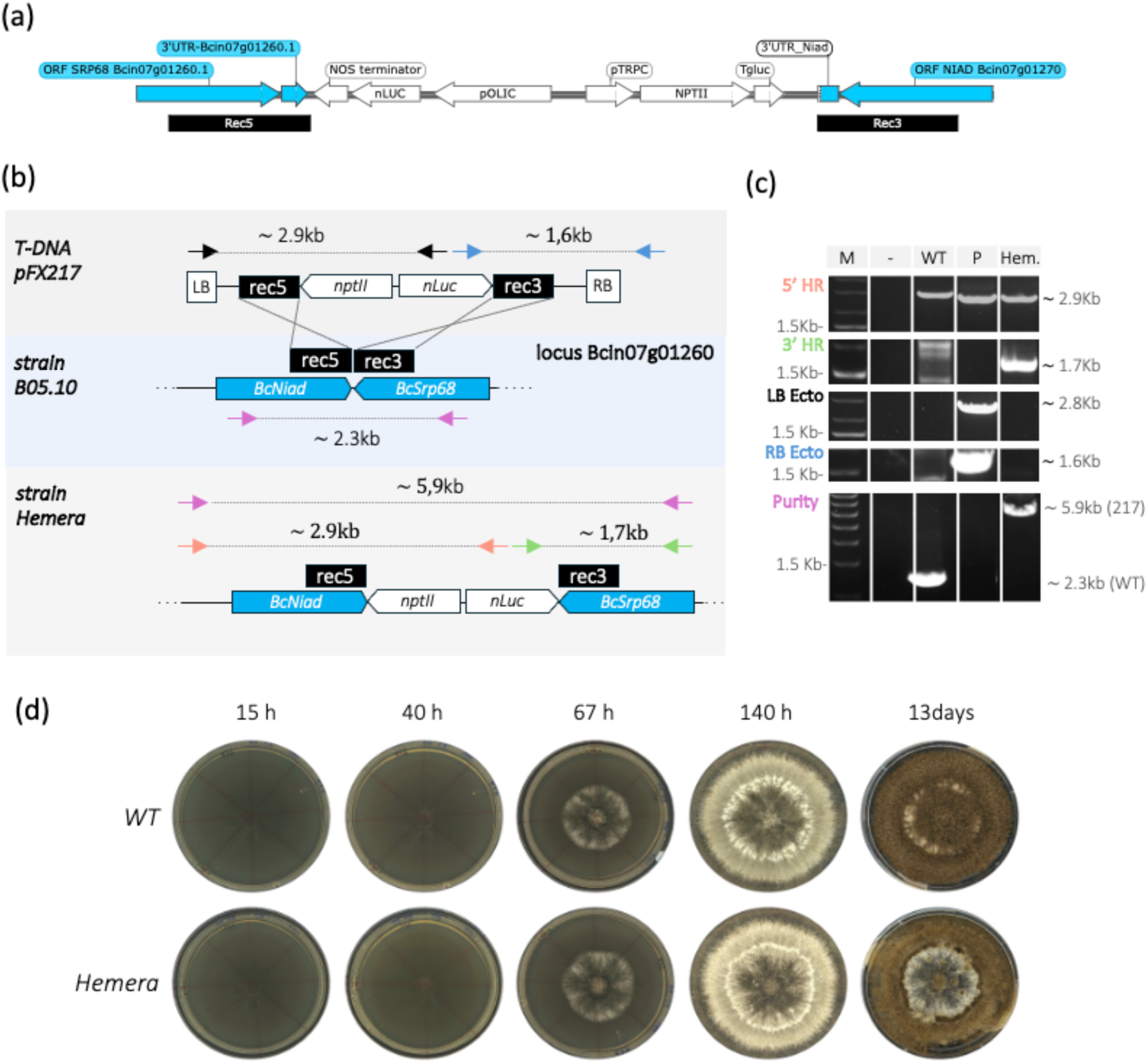
Generation and molecular and phenotypic characterization of the bioluminescent *Botrytis cinerea* reporter strain Hemera. **(a)** Schematic representation of the Hemera locus after targeted integration of the reporter construct into the intergenic region between *BcSRP68* and *BcNIAD*. The integrated cassette contains the constitutively expressed *nLuc* reporter and the *NPTII* selectable marker (neomycin resistant) together with their corresponding promoter and terminator sequences. **(b)** Strategy used for targeted integration and molecular verification. Top, organization of the T-DNA construct carrying the rec5 and rec3 homology arms, *nptII* and *nLuc*. Middle, parental B05.10 target locus. Bottom, resulting Hemera locus after homologous recombination. Colored arrows indicate the primer pairs used to verify the 5ʹ and 3ʹ integration junctions, detect potential ectopic T-DNA integration, and assess strain purity; expected PCR product sizes are indicated. **(c)** Molecular validation of Hemera by PCR. Assays were used to verify the 5ʹ and 3ʹ homologous recombination junctions (5ʹ HR and 3ʹ HR), test for left- and right-border ectopic integration (LB Ecto and RB Ecto), and assess strain purity after purification. M, molecular-weight marker; −, water control; WT, B05.10 genomic DNA; P, pFX217 plasmid; Hem., Hemera genomic DNA. A non-specific product slightly larger than the expected amplicon was detected in the WT lane of the 5ʹ HR assay. Five independent transformants were screened, and the transformant subsequently designated Hemera is shown. **(d)** Comparison of colony development and sporulation of parental B05.10 (WT) and Hemera on malt sporulation agar. Representative plates were imaged at 15, 40, 67 and 140 h after inoculation and after 13 days to assess sporulation. No obvious difference in colony development or sporulation was observed between the two strains under these conditions.

**Figure S2:**
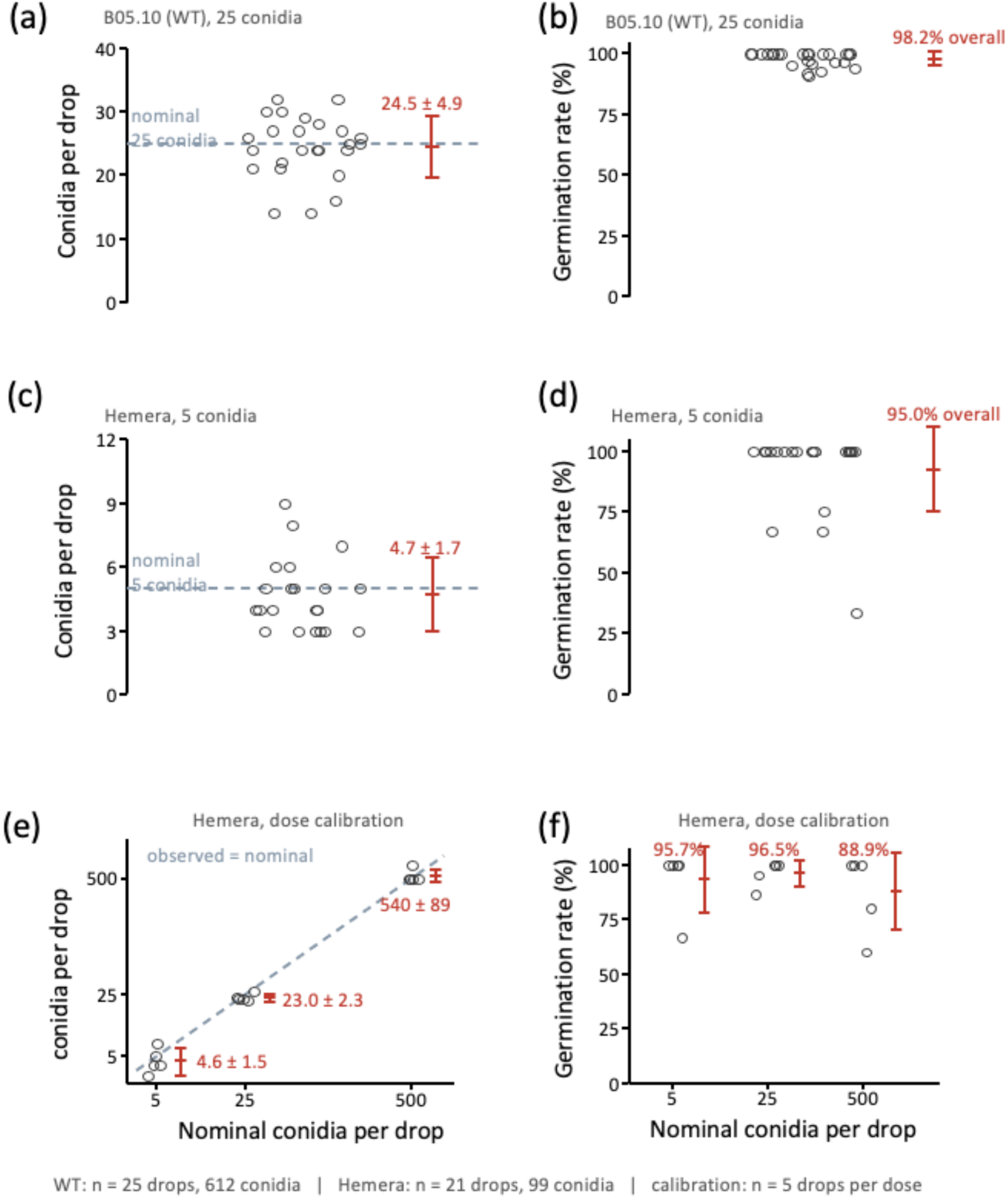
Calibration of the inoculum used in the seedling pathotest. Drops of 6.6 µL containing a defined number of conidia were deposited on 5 × 5 cm potato dextrose agar (PDA) plates and incubated overnight at 21 °C before germinated and non-germinated conidia were counted under a stereomicroscope. **(a, b)** *B. cinerea* B05.10 (WT), 25 conidia per drop (n = 25 drops, 612 conidia). **(c, d)** Strain Hemera, 5 conidia per drop (n = 21 drops, 99 conidia). **(e, f)** Strain Hemera at 5, 25 and 500 conidia per drop (n = 5 drops per dose); the 500-conidia condition was scored on a 1:100 dilution and the counts multiplied by 100. **(a, c)** Conidia recovered per drop; dashed line, nominal inoculum. **(b, d, f)** Germination rate per drop; percentages give the overall rate. **(e)** Observed versus nominal inoculum, logarithmic axes; dashed line, observed = nominal. Circles, individual drops; red bars, mean ± SD

**Figure S3.**
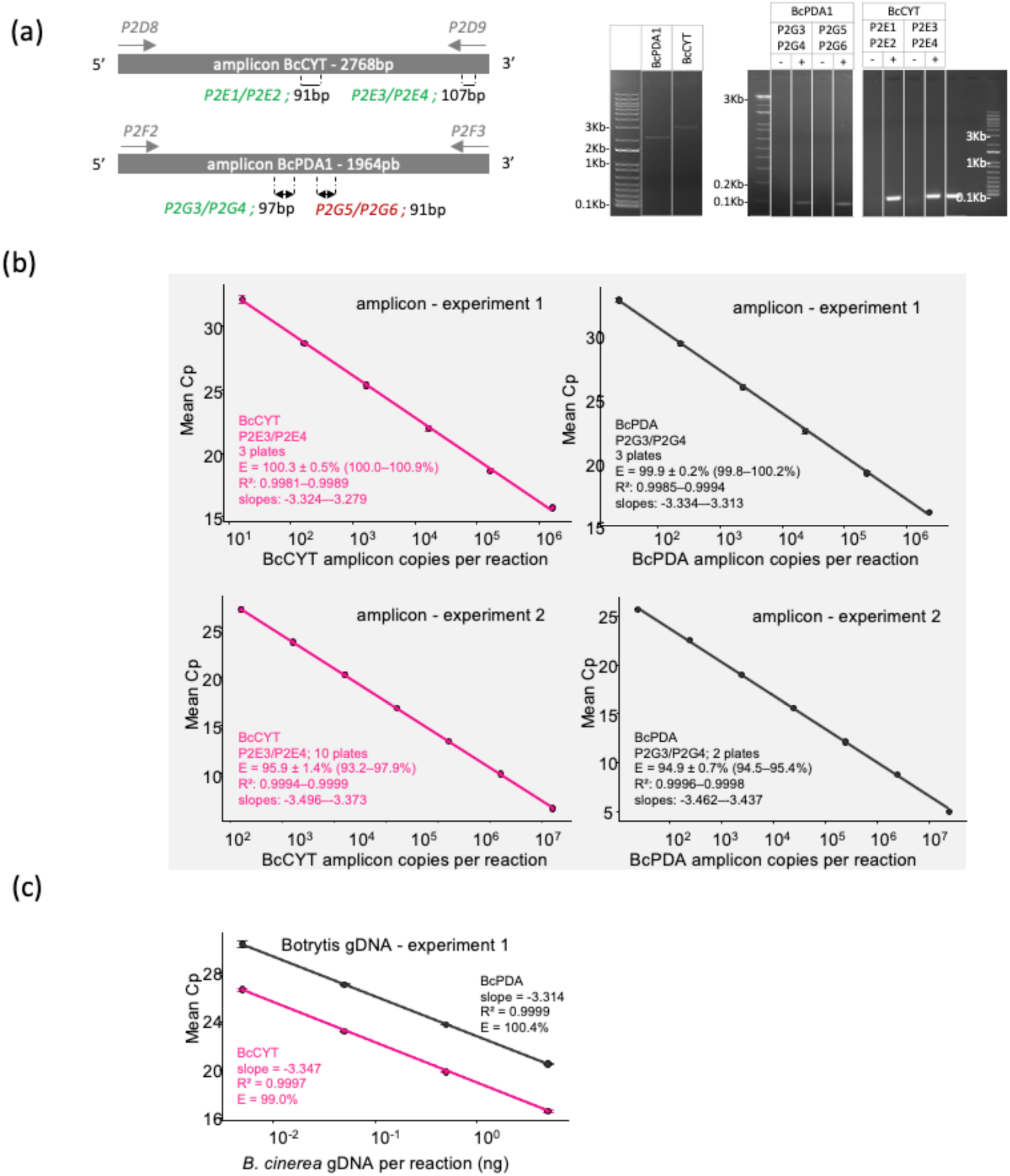
Mitochondrial and nuclear qPCR assays provide specific and highly efficient absolute quantification of *Botrytis cinerea* DNA. **(A)** Primer screening identifies specific BcCYT and BcPDA qPCR assays. Absolute qPCR assays were developed for a single-copy nuclear locus, BcPDA1 (Bcin07g01890), and a mitochondrial locus, BcCYT (BC1G-20024), in B. cinerea. Candidate primers were designed using NCBI Primer-BLAST. Two candidate primer pairs were generated for each locus using the following criteria: predicted (TM ∼ 60°C ; △ *2°C*), primer length (20–25nts), GC content (40–60%), 3’ G/C clamp, low predicted intra- or inter-primer complementarity, qPCR amplicon length (90–110 bp), and no predicted cross-amplification of A. thaliana. Positions of the primer pairs designed within the mitochondrial BcCYT and nuclear BcPDA amplicons are shown : in grey : template amplicons ; in green : qPCR primer pairs retained after validation ; in red : qPCR primers rejected (E>110%). Agarose-gel electrophoresis shows the purified template amplicons and the products generated by the candidate qPCR primer pairs. Negative controls are without Botrytis gDNA. **(B)** The selected assays display reproducible efficiencies across experiments. Standard curves were generated from serial dilutions of purified BcCYT and BcPDA amplicons in two independent experimental sessions (experiment 1 and experiment 2). Mean quantification cycle values are plotted against amplicon copies per reaction on a log10 scale. Lines indicate linear regressions. For the experiment 1, the highest-concentration standard was excluded because it consistently deviated from linearity for both assays. **(C)** BcCYT and BcPDA display high amplification efficiency and linear quantification over serial dilutions of *B. cinerea* genomic DNA. Standard curves were generated from serial dilutions of genomic DNA extracted from *in vitro*-grown *B. cinerea*. Mean quantification cycle values were plotted against the amount of fungal genomic DNA per reaction on a log10 scale. Linear regression parameters and calculated amplification efficiencies are indicated for each assay.

**Figure S4.**
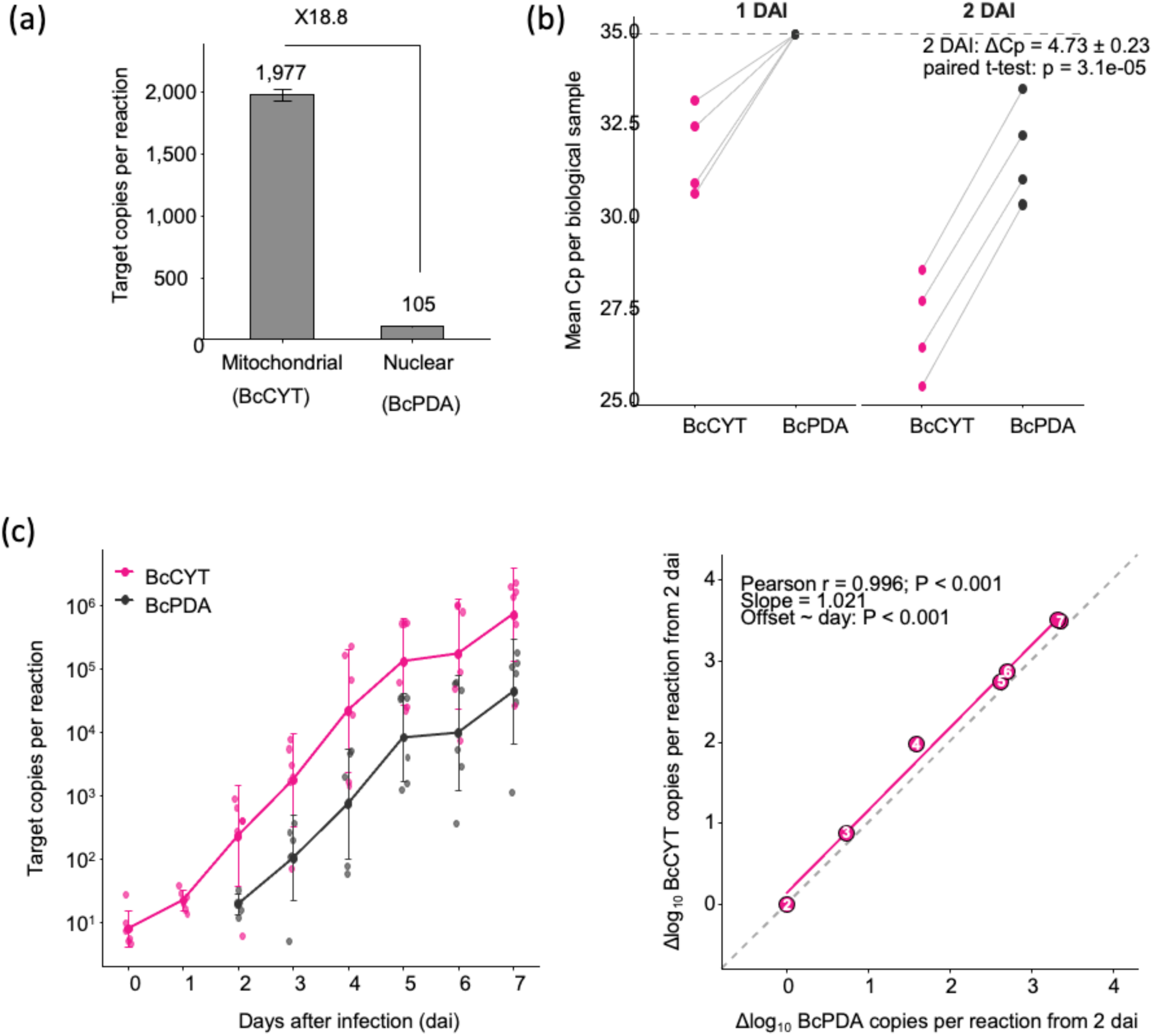
The mitochondrial BcCYT assay increases detection sensitivity while preserving the overall kinetics of fungal accumulation. **(a)** BcCYT is approximately 18.8-fold more abundant than the nuclear BcPDA target. Absolute target abundance was estimated from 0.05 ng of *B. cinerea* genomic DNA per reaction. **(b)** BcCYT improves early detection of fungal DNA in infected seedlings. BcCYT and BcPDA quantification was compared in infected seedlings at 1 and 2 days after inoculation (dai). Lines connect measurements obtained from the same biological sample. At 2 dai, BcCYT was detected at substantially lower Cp values than BcPDA, confirming the greater sensitivity of the mitochondrial assay at low fungal loads. **(c)** BcCYT and BcPDA capture the same overall growth dynamics despite a time-dependent mitochondrial-to-nuclear offset. Absolute target copy numbers were log10-transformed and are presented as mean ± SD, with individual biological replicates shown. The right-hand panel compares changes in the daily mean signal relative to 2 dai. Each labelled point represents one sampling day. The grey dashed line indicates perfect agreement and the solid fuchsia line the linear regression. The two assays showed strongly correlated temporal dynamics (r=0.996, P<0.001) and a regression slope close to unity (1.021). The mitochondrial-to-nuclear offset varied modestly over time (P<0.001). Thus, BcCYT increases detection sensitivity while preserving the overall temporal pattern of fungal accumulation. BcPDA values at 0 and 1 dai were omitted from the absolute kinetic analysis because the nuclear target was not quantitatively detected at these time points.

**Figure S5.**
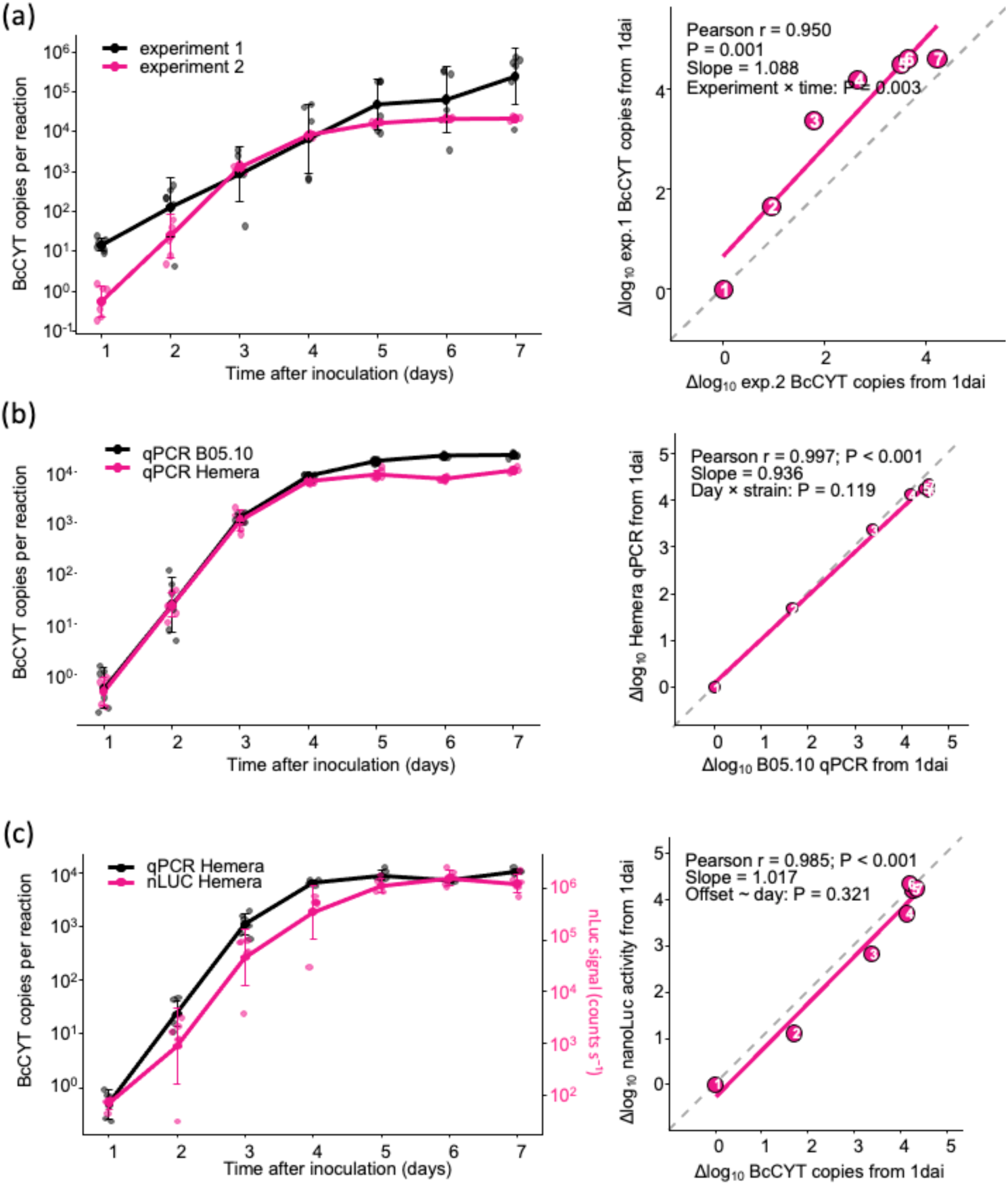
Independent pathotests vary in timing, whereas fungal colonization dynamics are highly reproducible within experiments across strains and quantitative methods. **(a)** Independent pathotests reproduce the same overall trajectory despite a shift in fungal expansion timing. B05.10 infection kinetics were compared between pathotests performed in experiment 1 and 2. The left panel shows absolute fungal DNA accumulation measured using the mitochondrial BcCYT assay. The right panel compares changes in mean log10- transformed BcCYT copy numbers relative to 1 dai. The two experiments displayed strongly correlated dynamics (r=0.950, P=0.001) and a regression slope close to unity (1.088). However, the significant experiment × time interaction (P=0.003) indicates a moderate shift in the magnitude or timing of fungal expansion between pathotests. **(b)** B05.10 and Hemera display nearly identical fungal colonization dynamics within the same pathotest. Changes in mean log10-transformed BcCYT copy numbers were calculated relative to 1 dai. The two strains showed highly similar temporal dynamics (r=0.997, P<0.001; slope = 0.936), with no significant strain × day interaction (P=0.119). **(C)** qPCR and bioluminescence provide highly concordant measurements of fungal accumulation. Fungal accumulation in the Hemera strain was quantified using BcCYT qPCR and nanoLuc enzymatic activity. Changes in both signals were calculated relative to 1 dai. The two measurements showed strongly concordant dynamics (r=0.985, P<0.001; slope = 1.017), and their offset did not vary significantly over time (P=0.321). In all kinetic plots, points represent biological replicates, whereas lines and error bars represent mean ± SD. Numbers within the concordance plots indicate sampling days. Dashed grey lines represent perfect agreement, and solid fuchsia lines indicate linear regressions. Together, these analyses show that the exact timing of fungal expansion varies between independent pathotests, whereas comparisons performed within the same experiment remain highly reproducible across strains and quantitative approaches.

**Figure S6.**
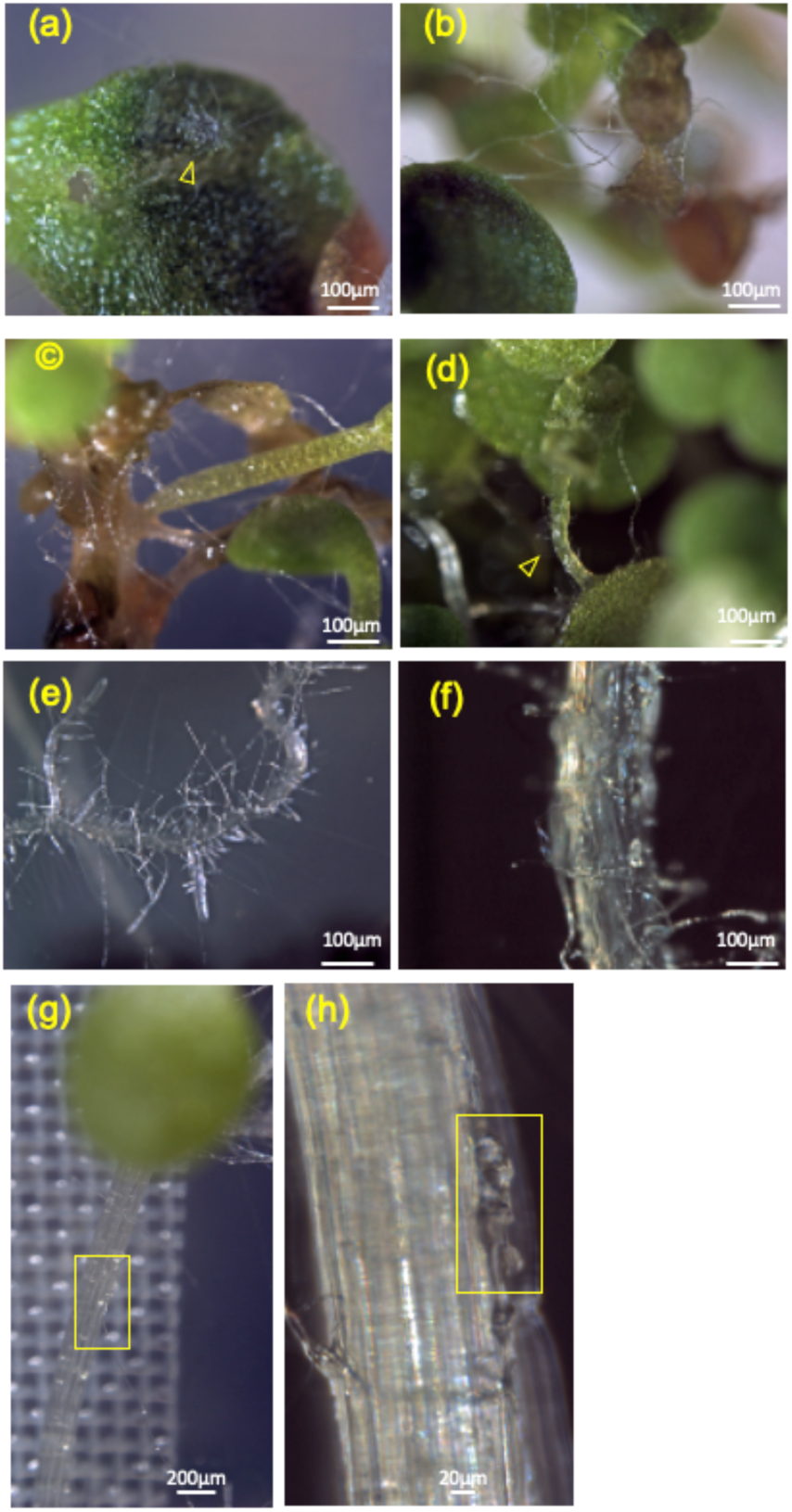
Representative macroscopic and microscopic features of *Botrytis cinerea* colonization of *Arabidopsis* seedlings. **(a)** Extensive surface-associated fungal growth on cotyledon (4 dai); the arrow indicates an infection cushion. **(b,c)** Seedlings illustrating advanced disease development at 5 dai and extensive tissue necrosis. **(d)** Cotyledon petiole (5 dai) displaying a translucent, glassy appearance, with fungal hyphae visible at the tissue surface. **(e,f)** Root colonization (3 dai), showing fungal hyphae extending along the root surface. **(g,h)** Infected hypocotyl (3dai). The yellow rectangle in panel g indicates the region imaged in h. The yellow rectangle in h is the region imaged by confocal microscopy shown in figure 5 panel c. Scale bars are indicated in the corresponding panels.

**Figure S7.**
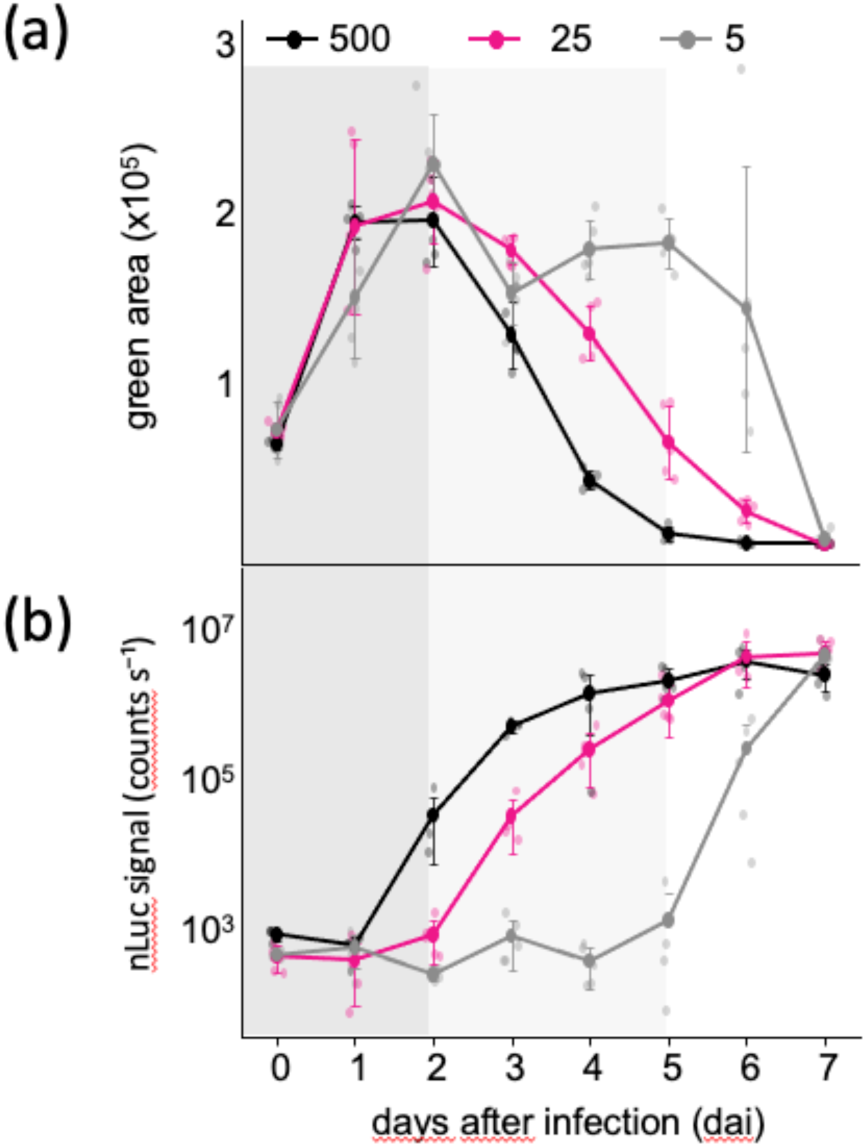
Pathotest variability underlying T10⁵ alignment. (a) Individual bioluminescence trajectories of all pathotest inoculated with 500, 25 or 5 spores. The dashed horizontal line indicates the T10⁵ threshold used to define fungal establishment. (b) The same trajectories after independently aligning each biological replicate to its own T10⁵ establishment time. Alignment removes differences in establishment timing and reveals the reproducibility of the expansion phase across inoculum densities.

**Figure S8.**
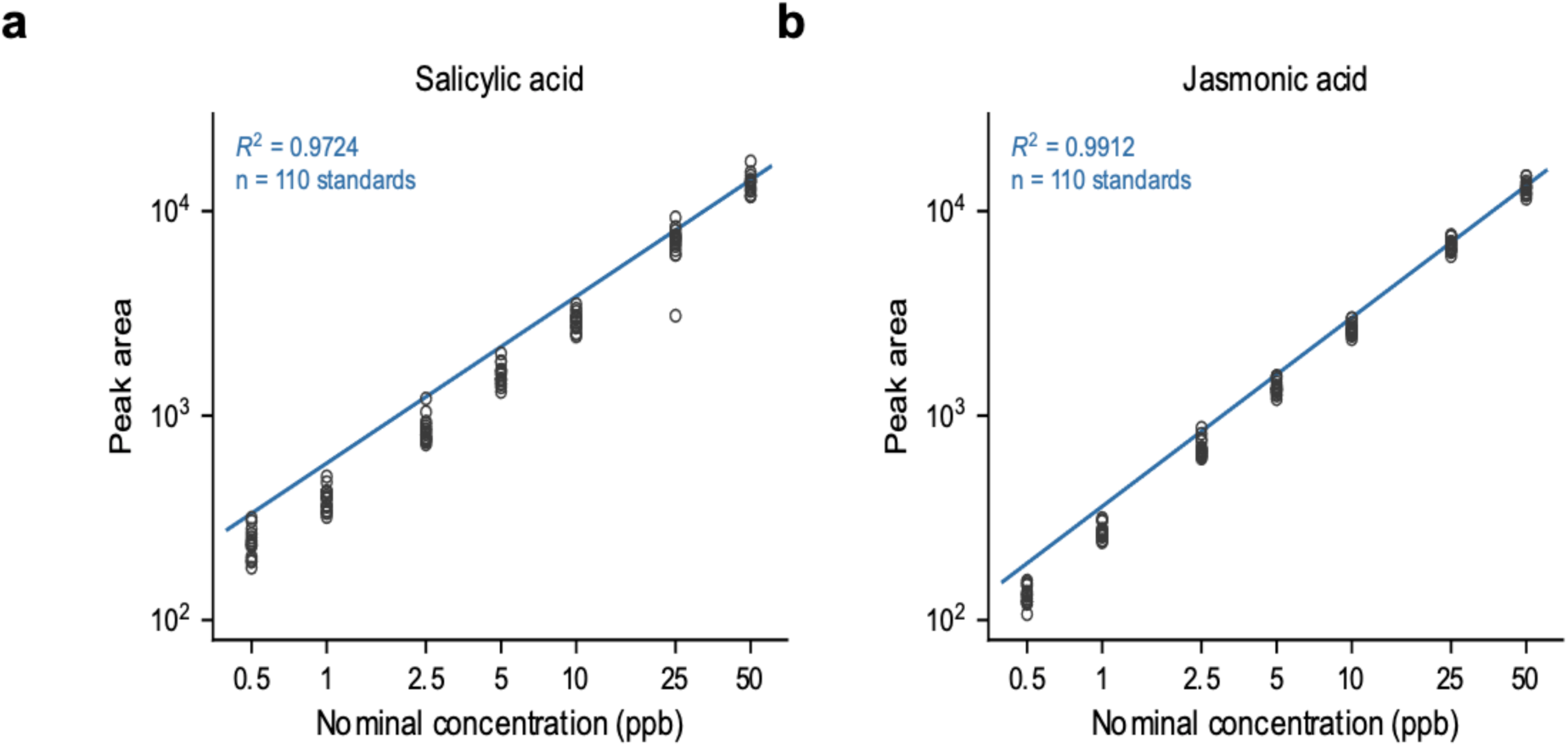
: Calibration of the LC-MS/MS quantification of salicylic and jasmonic acid. Standards containing both analytes at 0.5, 1, 2.5, 5, 10, 25 and 50 ppb were injected at the start and at the end of each of the eight analytical sequences of the time-course experiment. (a) Salicylic acid. (b) Jasmonic acid. Each circle is one standard injection and the line is the linear regression fitted to all injections, on logarithmic axes (n = 110 injections per analyte; two injections per analyte were excluded after instrument failure).

**Figure S9.**
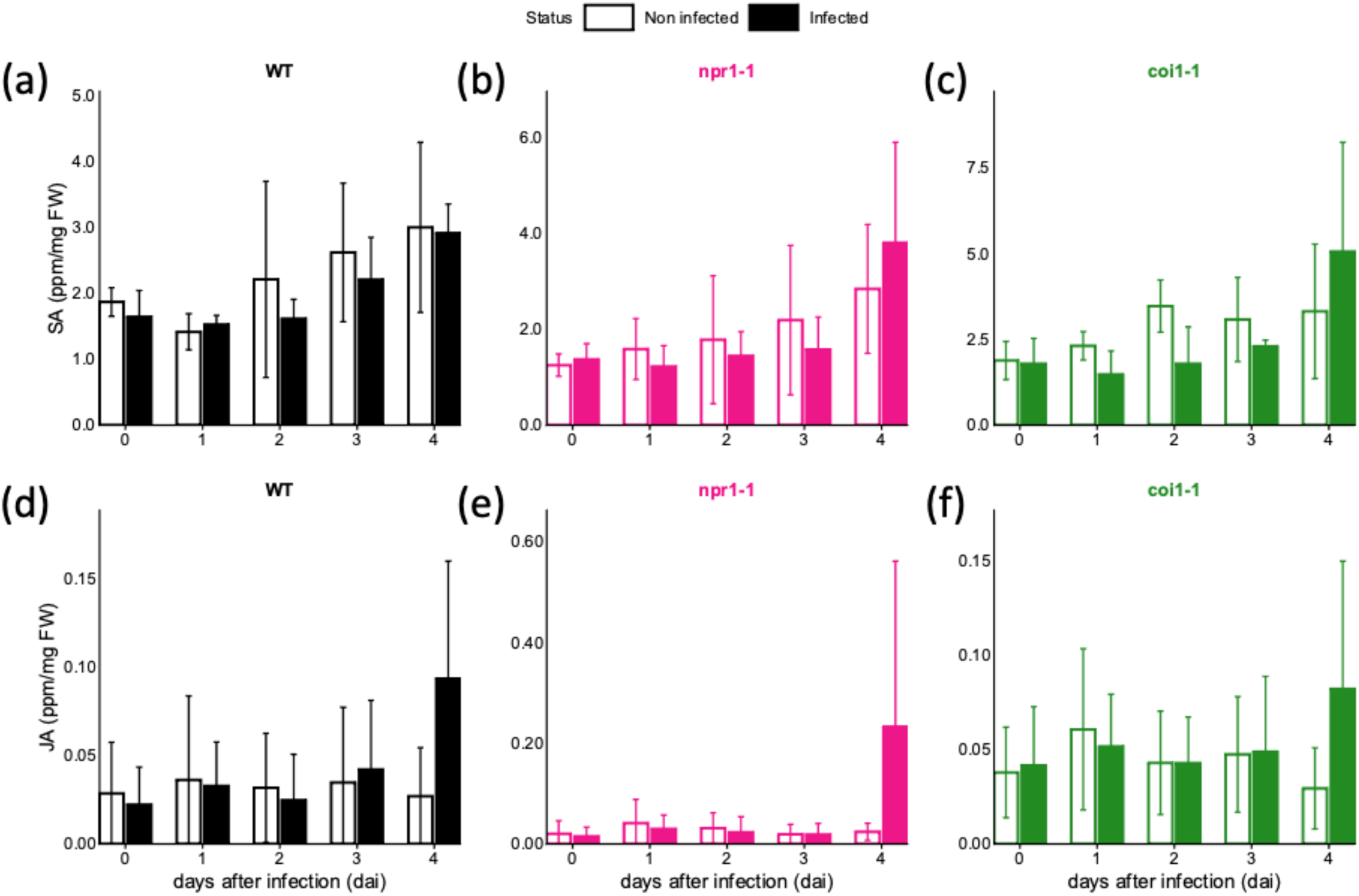
Hormone levels remain comparable between infected and non-infected seedlings during the asymptomatic phase across all three genotypes. Salicylic acid (SA, top row) and jasmonic acid (JA, bottom row) contents were quantified by LC–MS/MS in wild-type (WT), *npr1-1* and *coi1-1* seedlings from 0 to 4 days after inoculation (dai) - (n=6). For each genotype, hormone levels in non-infected and *Botrytis cinerea*-infected seedlings are shown as mean ± SD. Statistical comparisons between infected and non-infected samples were performed independently at each time point using two-sided unpaired Wilcoxon tests, followed by Benjamini–Hochberg correction across the five time points within each genotype and hormone. Only significant adjusted comparisons would be displayed on the plots. No statistically significant differences were detected before 5 dai, indicating that neither the SA nor the JA pathway is robustly activated during the early asymptomatic phase of infection.

**Figure S10.**
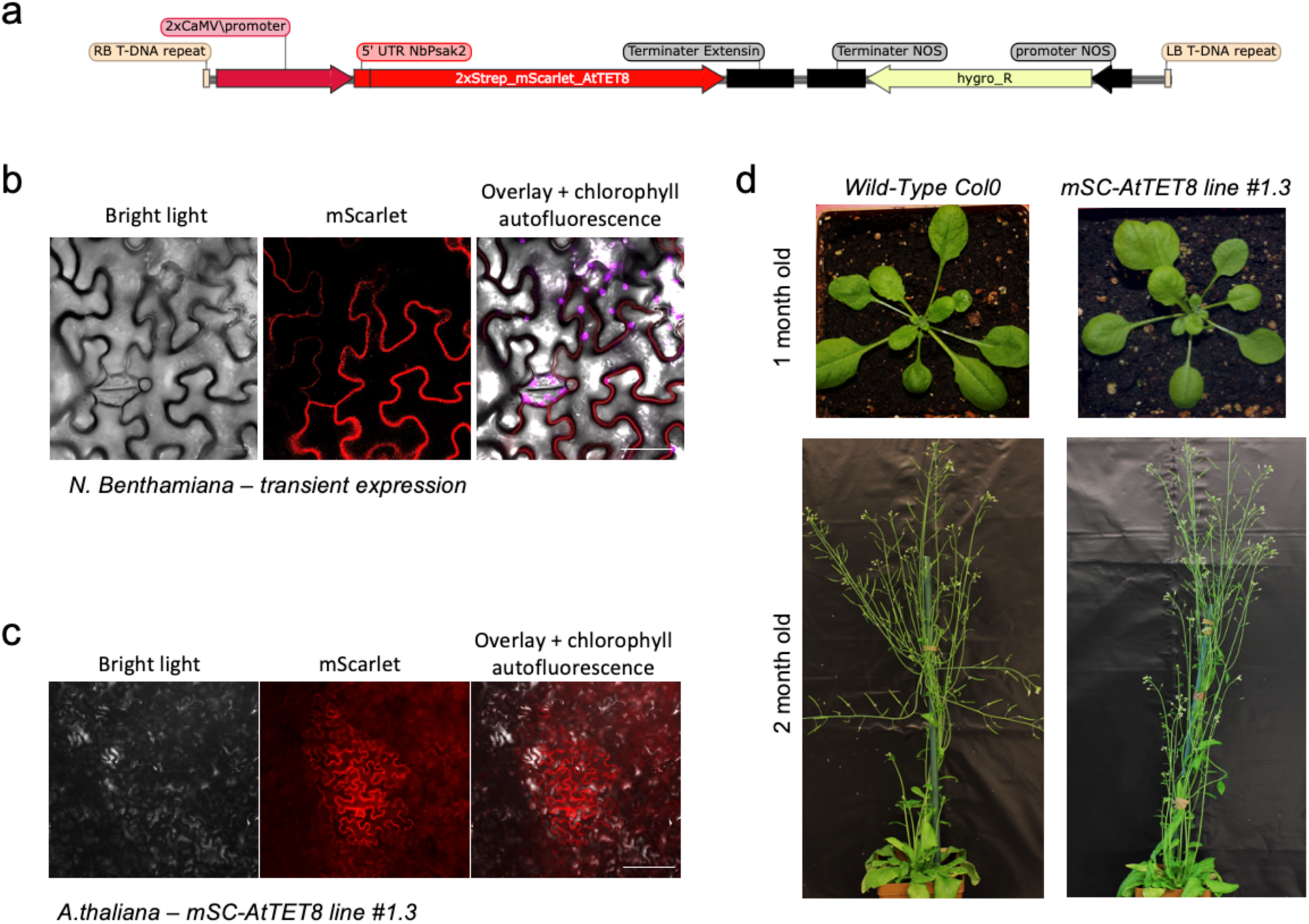
Generation and phenotypic validation of the *Arabidopsis thaliana* mScarlet-AtTET8 reporter line. (a) Schematic representation of the 2×35S:mScarlet-AtTET8 construct used for plant transformation. (b) Transient expression of mScarlet-AtTET8 in *Nicotiana benthamiana* epidermal cells, showing the expected cell-peripheral fluorescence pattern. (c) mScarlet-AtTET8 fluorescence in epidermal cells of the stable *A. thaliana* line #1.3. Bright-field, mScarlet fluorescence and overlays including chlorophyll autofluorescence are shown in (b) and (c). Scale bars, 20 µm. (d) Representative morphology of wild-type Col-0 and homozygous mScarlet-AtTET8 line #1.3 plants at 1 and 2 months of age, showing no obvious alteration in overall plant development.

**Figure S11.**
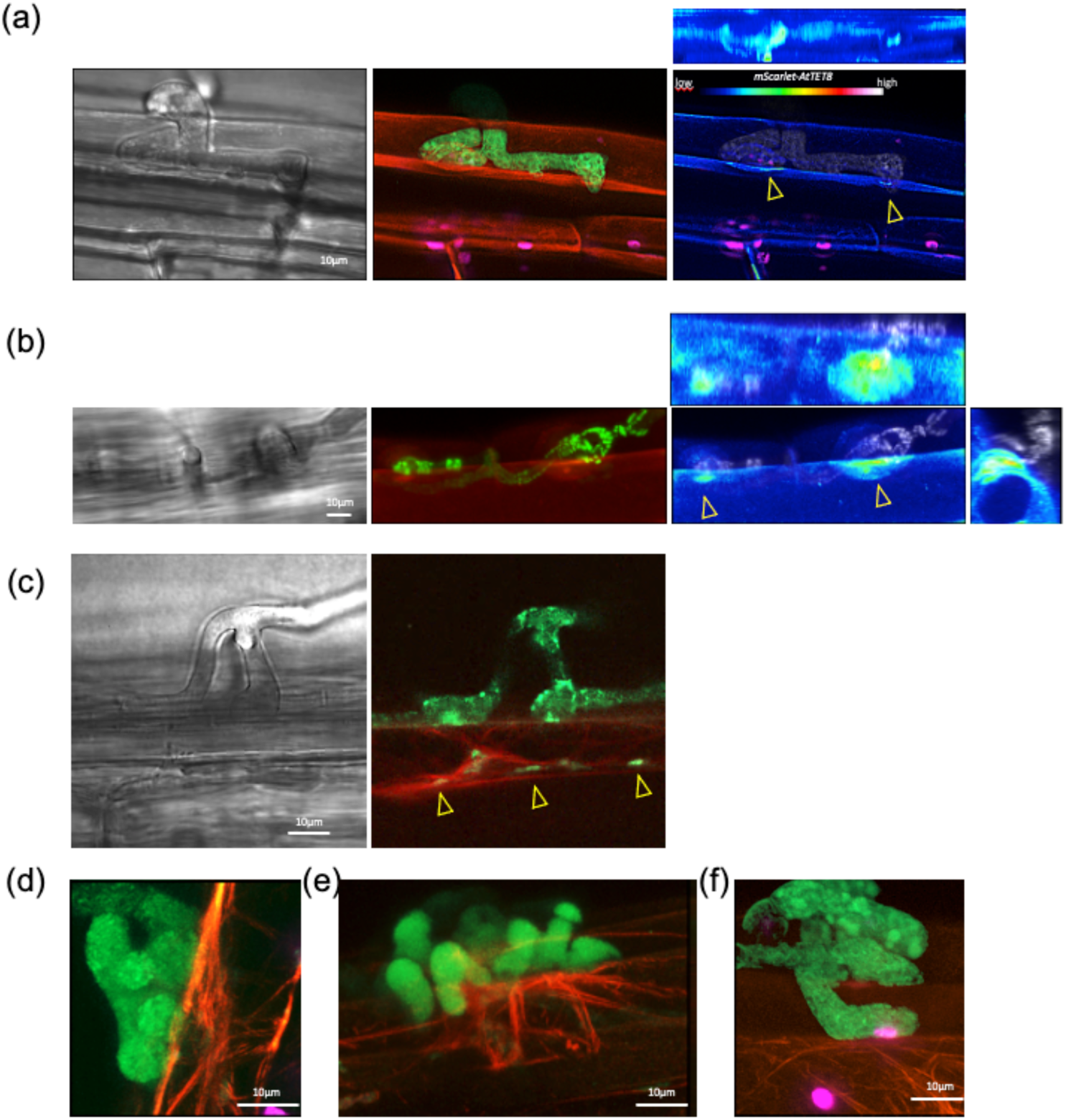
Additional examples of localized TET8 enrichment and cortical actin responses at the *Botrytis–Arabidopsis* interface. Live-cell confocal imaging was performed on *Arabidopsis thaliana* hypocotyl tissues infected with a *Botrytis cinerea* strain expressing PLS1- mNeonGreen. Depending on the panel, host cells expressed either mScarlet-TET8 or a LifeAct reporter. **(a)** Example of a fungal invasion event associated with local host TET8 accumulation. *B. cinerea* penetrates a host cell, and arrowheads indicate localized enrichment of the TET8 signal at the host–fungus interface. Orthogonal views illustrate the spatial relationship between the fungal structure and the TET8- enriched host domain. **(b)** Example of endophytic fungal progression between host cells associated with local TET8 enrichment. Arrowheads indicate localized accumulation of TET8 at the host interface facing the fungal structure. The higher-magnification view highlights the contact site. **(c)** Example of a differentiated fungal structure showing detectable PLS1-mNeonGreen signal within the host cell volume, in close proximity to the host cortical actin network. In this interaction, no obvious actin polarization toward the fungal structure is detected. **(d,e)** Additional examples of cortical actin polarization toward differentiated fungal structures. Actin bundles locally converge toward the fungus– host interface. **(f)** Example of a fungal contact site in which close association with the host cell is observed, but no obvious polarization of the cortical actin network is detected. For selected panels, mScarlet-TET8 fluorescence intensity is displayed using a rainbow lookup table (LUT) to facilitate visualization of local signal enrichment. Scale bars are indicated in the corresponding panels.

## Acknowledgements

We thank Martin Donnadieu (MAP, Lyon, France) from our plant facility. We are grateful to Marie-Cécile Caillaud (RDP, ENS de Lyon, France) for kindly providing the *Arabidopsis thaliana* Lifeact-tdTomato line. We thank Marcel Calleja (Bayer, Lyon, France) for facilitating safe sample collection on non-working days. We also thank Rachel Baltz (Bayer, Lyon, France) for her support throughout the project. We thank Mathilde Fagard (IJPB, Versailles, France), Thomas Guillemette (IRHS, Angers, France), Ludovic Vial (LEM, Lyon, France), Alexandre Soulard (MAP, Lyon, France) and Daniel Muller (LEM, Lyon, France) for valuable discussions and feedback.

## CRedit authorship contribution statement

Conceptualization : F.-X.G., C.Bo, ; Supervision : F.-X.G ; Investigation : I.T., P.F., A.P., F.-X.G ; Methodology : I.T., C.Bo, F.-X.G. ; Data curation : I.T., F.-X.G. ; Formal analysis : I.T., F.-X.G., C.Bo ; Writing – original draft - review and editing : I.T., C.Bo, F.-X.G. ; Writing –review and editing : N.P., M.C., C.Br.,

